# Increased dispersal explains increasing local diversity with global biodiversity declines

**DOI:** 10.1101/2023.06.09.544194

**Authors:** Brennen Fagan, Jon W. Pitchford, Susan Stepney, Chris D Thomas

## Abstract

The narrative of biodiversity decline in response to human impacts is overly simplistic because different biodiversity metrics show different trajectories at different spatial scales. It is also debated whether human-caused biodiversity changes lead to subsequent, accelerating change (cascades) in ecological communities, or alternatively build increasingly robust community networks with decreasing extinction rates and reduced invasibility. Mechanistic approaches are needed that simultaneously reconcile different metrics of biodiversity change, and explore the robustness of communities to further change. We develop a trophically-structured, mainland-archipelago metacommunity model of community assembly. Varying the parameters across model simulations shows that local alpha diversity (the number of species per island) and regional gamma diversity (the total number of species in the archipelago) depend on both the rate of extirpation per island and on the rate of dispersal between islands within the archipelago. In particular, local diversity increases with increased dispersal and heterogeneity between islands, but regional diversity declines because the islands become biotically similar and local one-island and few-island species are excluded (homogenisation, or reduced beta diversity). This mirrors changes observed empirically: real islands have gained species (increased local and island-scale community diversity) with increased human-assisted transfers of species, but global diversity has declined with the loss of endemic species. However, biological invasions may be self-limiting. High-dispersal, high local-diversity model communities become resistant to subsequent invasions, generating robust species-community networks unless dispersal is extremely high. A mixed-up world is likely to lose many species, but the resulting ecological communities may nonetheless be relatively robust.

**Significance Statement:** Biodiversity is commonly regarded as threatened due to human impacts, but biodiversity metrics at different scales produce contradictory results. A framework is needed that can reproduce and connect these results across scales and address whether biodiversity change will inexorably accelerate following perturbation or become self-limiting as new ecological communities form. We address this challenge by constructing size-structured model communities using a mainland/island paradigm and tracking diversity at different scales. Our simulations reproduce the literature’s discrepancy across scales and provide new insight. Ecological communities (islands) gain species with increasing (human-assisted) dispersal, but global diversity declines with the consequent loss of endemic species. Communities also become less invasible as dispersal increases, suggesting that human-mediated dispersal favours robust communities that resist subsequent change.

---

**M**ultiple human-associated pressures are driving changes to biological diversity worldwide, leading to both academic and popular concern over the impending ‘biodiversity crisis’ and possible ‘Sixth Mass Extinction’ (1, 2). In particular, there is great concern that the number of species per unit area (alpha diversity) is declining due to land use changes and intensification (3), that species-level rates of endangerment and extinction are increasing (reducing the total number of species, or gamma diversity) (4, 5), and that land use change combined with the transport of species between different geographic locations is resulting in biological homogenisation (reduced beta diversity), as well as causing the extinction of localised species whose ranges have been invaded (6–8). Taken together, these works strongly support the contention that ‘biodiversity’ is endangered, however it is measured.

In contrast, other studies suggest that alpha, beta and gamma diversity trajectories depend on the spatial scale and locations studied (9). Many studies have found that local alpha diversity is increasing in some locations while declining in others, with average alpha remaining similar, or possibly even increasing slightly (10–12). At a larger scale, regional diversity has mainly increased despite reduced global (gamma) diversity: the number of plant species on oceanic islands has increased with plant invasions despite the extinctions of many island-endemic species (13), the number of plant species per country or state is increasing despite regional losses (14), and the number of mammal species has increased in most European countries over the last 8,000 years despite many species having become extinct globally (15). And, in contrast to studies reporting homogenisation at some scales (above), land use changes may have increased beta diversity at many spatial scales for most of the last millennium because different species are associated with different ‘natural’ and human-modified ecosystem types (16). This flurry of studies generating different directions and strengths of alpha, beta and gamma diversity change in different spatial, temporal and geographical contexts has led to confusion and disputes in the literature, undermining the development of consensus measures to manage biodiversity (e.g., 17–25).

Further confusion relates to the extent to which these derived ecosystems and ecological communities become more or less resistant to further biological change through time. Invasion biology identifies the potential for community ‘cascades’ and ‘collapse’ in which community turnover (loss of some species, invasion of others) begets further turnover (26, 27). This perspective holds that the initial changes to community food webs lead to successive changes, resulting in further losses and increased potential for subsequent invasions as community networks reorganise. Adaptive community dynamics (28, 29), in contrast, holds that changes to the identities and relative abundances of species in a given location should generally increase the robustness of certain community properties as they adjust to both the (new) biological and physical environments, potentially increasing their resistance to further invasion. Thus, the dynamic processes of community reorganisation as well as the outcomes (scales of diversity change) are debated.

Community ecology has already seen that altered rates of movement (inter-patch mixing) drive local diversity and homogenisation simultaneously (30, 31), as in Figure 1. We further develop the theory of inter-patch mixing, henceforth dispersal, and how dispersal affects not only the observed patterns of biodiversity across scales but also ongoing and future changes to communities by considering temporal turnover and invasibility. Using a mainland-archipelago model, we find that changing the rate of dispersal of organisms between heterogeneous islands within the archipelago is qualitatively sufficient to generate the variety of contrasting biodiversity trends observed in nature; while immigration events and local extirpations do generate ongoing community dynamics, high dispersal generally produces communities that are robust to further invasion. Simple dynamics in the model can generate multiple patterns in the simulated communities in space and time, helping to resolve several apparent paradoxes in assessments of different metrics of biodiversity change.

**Fig. 1.**
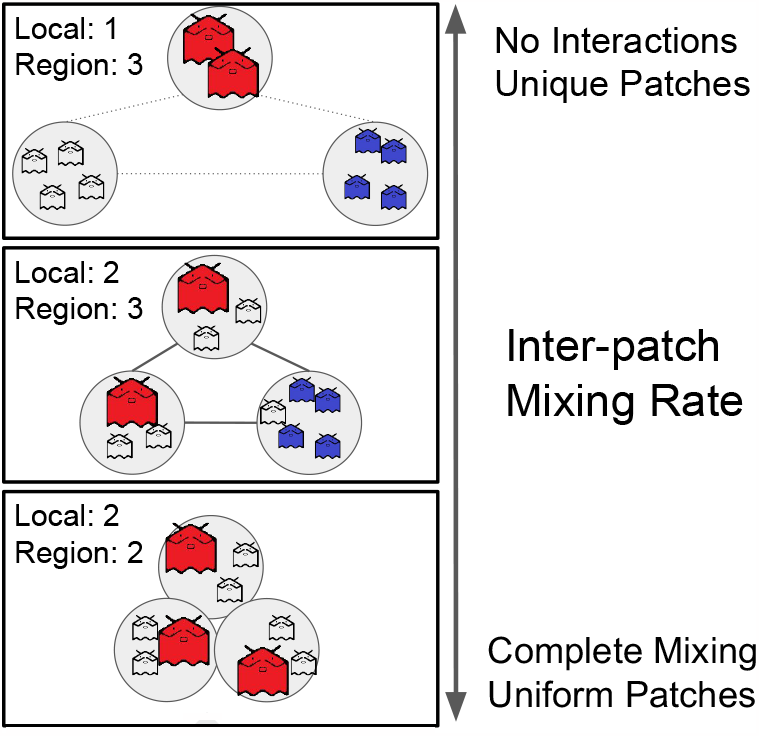
Hypothesised relationships of local and regional diversity metrics with dispersal. In communities with negligible dispersal (top), local communities (alpha) support few species (here, 1 critter species per patch/island), but different species (high beta), creating high regional gamma diversity (here, 3 critter species in total). High dispersal (bottom) generates richer local communities (here, 2 per patch) but the communities are the same as one another due to species mixing, and hence total regional diversity is only 2 species. At intermediate mixing (middle panel), various outcomes are possible, here showing local communities as diverse as complete mixing, and regional richness as diverse as no mixing (one variant of the intermediate dispersal hypothesis).

## Model Description

Community ecology is no stranger to contradictory or counter-intuitive results and a mismatch between data and theory, including May’s seminal result that complex communities are unlikely to be stable (32), a view not necessarily held by empirical ecologists (33). The ensuing debates have required ecologists and mathematicians to understand which mechanisms and assumptions need to be changed in order for models to usefully reflect reality, leading to valuable developments in theory (33) and in the interpretation of empirical data (34–36). Both theoretical and empirical studies consistently point to the importance of size- and trophic structure in shaping local communities, and to the role of community assembly (the sequential local rearrangement of communities based on arrivals – long distance dispersal – and extinctions) in forming functional ecosystems (34, 36–40). The importance of both the structure of communities and space (affecting dispersal and interactions) have been emphasised in recent studies (36, 41–43). Increased dispersal can potentially complex biodiversity trends (e.g., increased alpha by enlarging the effective immigrant pool, but reduced beta via homogenisation of communities) including the maximisation of local diversity at intermediate dispersal (31), but more complex patterns may emerge (44, 45). Our simulations that incorporate both dispersal and local community dynamics generate peak local diversities at intermediate dispersal when environments differ between patches (with the peak height increasing with the heterogeneity between patches), and opposing biodiversity trends when measured using different metrics.

Our model is an extension of community assembly, which mirrors island biogeography, and our analyses include interpatch mixing. In a community assembly method, one seeks stable and complex systems by constructing a regional pool (a metaphorical mainland) from which species can migrate to initially empty patches (islands) via small initial populations. This method is known to successfully create systems of moderate size that can be resistant to further invasion from the pool in a theoretically robust and consistent manner (37, 38). Thus, we have two major components to the model: community interactions and movement. We summarise the model in Figure 2 and briefly touch on its features here; we describe them further in the Materials and Methods and Supplemental Information.

**Fig. 2.**
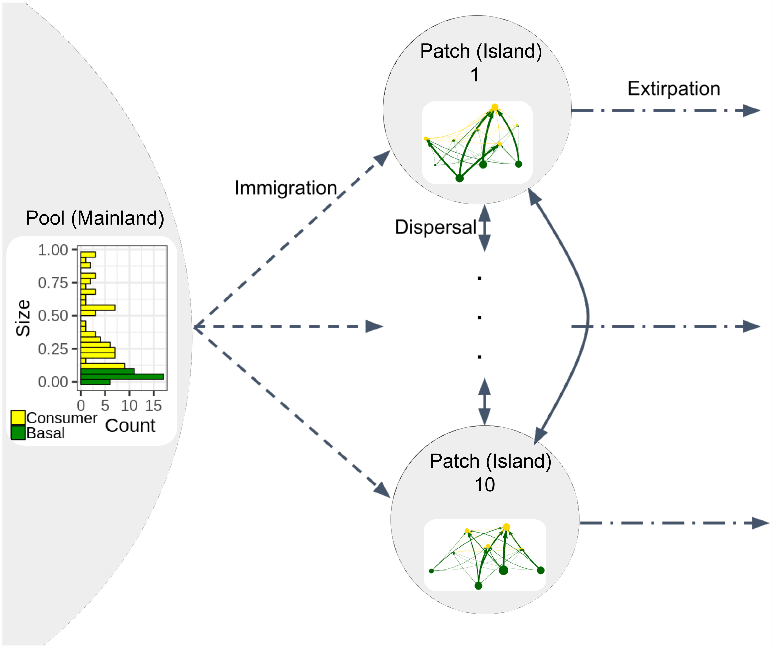
Model set-up at multiple scales. Our model begins with a fixed size-structured pool of species (left) with size-structured predation relations (histogram of size (vertical axis), with green for basal species and yellow for consumer species) and a fixed number of patches. The patches are allowed to differ abiotically from each other, slightly changing the strengths of interspecific interactions. Throughout the simulation, immigration can occur in which a sub-population of a species is drawn randomly from the pool and ‘tries’ to colonise a randomly selected patch (dashed lines). The sub-population successfully establishes if its per-capita growth rate is positive for the given patch without dispersal. Species that successfully establish are then subject to community dynamics within their patch food webs (with the size of each circle corresponding to a species’ biomass and thickness of connecting lines representing the amount of flow), and can disperse in proportion to their abundance (along the double-headed arrows) to neighbouring patches (here patches are connected as a ring). Dot-dash lines from each patch indicate both deterministic extirpations (arising from community interactions) and also random, neutral extirpation events. The histogram here represents the pool used for the simulations in Figure 3, and the patch networks are representatives from a single time step of the no dispersal simulation. Network visualisation uses foodwebviz (46).

The community interactions are created by following the process of Law and Morton (37). Sizes are fixed for the species pool and then used to determine all species properties and the trophically structured interaction matrices. Whilst size structured (some species function as autotrophs and heterotrophs exploit autotrophs and other heterotrophic species within a size range smaller than themselves), in principle similar outcomes are expected for any kind of consumer-consumed interaction network. Community interactions are normally assumed to take place in between arrivals – time scale separation – but we allow them to proceed continuously in parallel to facilitate the study of dispersal. We also then allow interaction strengths to vary between patches to reflect how environmental conditions may influence interspecific interactions.

In the model, we differentiate species movement between long-range pool–patch immigration and short-range patch– patch dispersal. The simulations start with no species present within a network of similar but not identical patches (the conceptual archipelago, see Materials and Methods) and the fixed species pool (conceptual mainland). Species immigrate to the archipelago (drawn at random from the pool, a neutral process) by sending a sub-population into a single patch (conceptual island; dashed arrows in Figure 2). The rate of immigration can be varied, although we focus here on one rate of immigration which is similar to the rate of community interactions (but consider sensitivity to this variable). If immigrants of a new species establish a population, it results in colonisation. Hence it is possible to evaluate the robustness of communities to subsequent colonisation (fraction of possible immigration events resulting in colonisation; conceptually the invasibility of communities). At the end of a simulation, we test the capacity of each community (considering each patch separately) to resist invasion by deliberately and separately introducing each species (not already present) from the species pool.

As communities establish in each patch, individuals from those communities are (neutrally) mixed with individuals (‘dispersal’) from other communities (see Materials and Methods for details, vertical arrows in Figure 2), varying from zero mixing to complete mixing. The patches – in our case 10 – are arranged in a ring, so communities are up to five dispersal steps away from one another. Thus, our island archipelagos operate as metacommunities. When discussing variation in dispersal, we focus here on this between-patch mixing rate. We refer to pool-to-patch movement of species as ‘immigration’ and patch-to-patch movement as ‘dispersal’ (Figure 2).

Species extirpations (loss from a single patch) and extinctions (loss from the entire archipelago, but not from the pool so re-invasion is possible) take place (dash-dotted arrows in Figure 2). These are commonly driven by community dynamics, but we also include a stochastic element, applied at the patch level as is commonly adopted in stochastic metapopulation and metacommunity models to mimic local disasters including potentially human-mediated ones (see Supplemental Information for the no stochastic extirpation case). Thus, we include both deterministic (via community network dynamics) and stochastic components of species removals from communities.

Our simulation results allow us to explore the interactions between an ecologically justified model, dispersal, heterogeneous environments, and the pool-patch structure. These simple modifications to community assembly are sufficient to recreate the observed phenomenon of per island alpha diversity increases and archipelago-scale gamma diversity declines. They are also sufficient to allow us to explore the impacts of our parameter choices on the spatial dissimilarity between patches at an instant in time (beta diversity calculated using the Jaccard dissimilarity index, see Materials and Methods), the temporal turnover (Jaccard dissimilarity index through time within a patch) and the final invasibility of the system (see Materials and Methods), which characterise whether our communities stabilise through time within islands and across the entire archipelago.

## Results

### Example Runs

Figure 3 illustrates three runs with the same pool of species, same set of patch environments, and precisely the same history of immigration and extirpation events (which does not guarantee the same history of successful establishments if the communities differ, e.g., due to dispersal). The only difference in parameters is the change in dispersal from no dispersal between patches (zero inter-patch mixing case), to ‘medium’ and full dispersal cases, in which some or nearly all of the abundance on a patch can have moved to an adjacent patch in a single time unit respectively (these example pools, environments, and histories have been selected at random). The no-dispersal example has many transient species (short horizontal dashes in the upper panels of Figure 3), most of which occur in only a small number of patches at any one time (yellow), whereas the high dispersal example shows a smaller number of species present in most patches (purple) for most of the simulation run (long dashes); medium dispersal is intermediate.

**Fig. 3.**
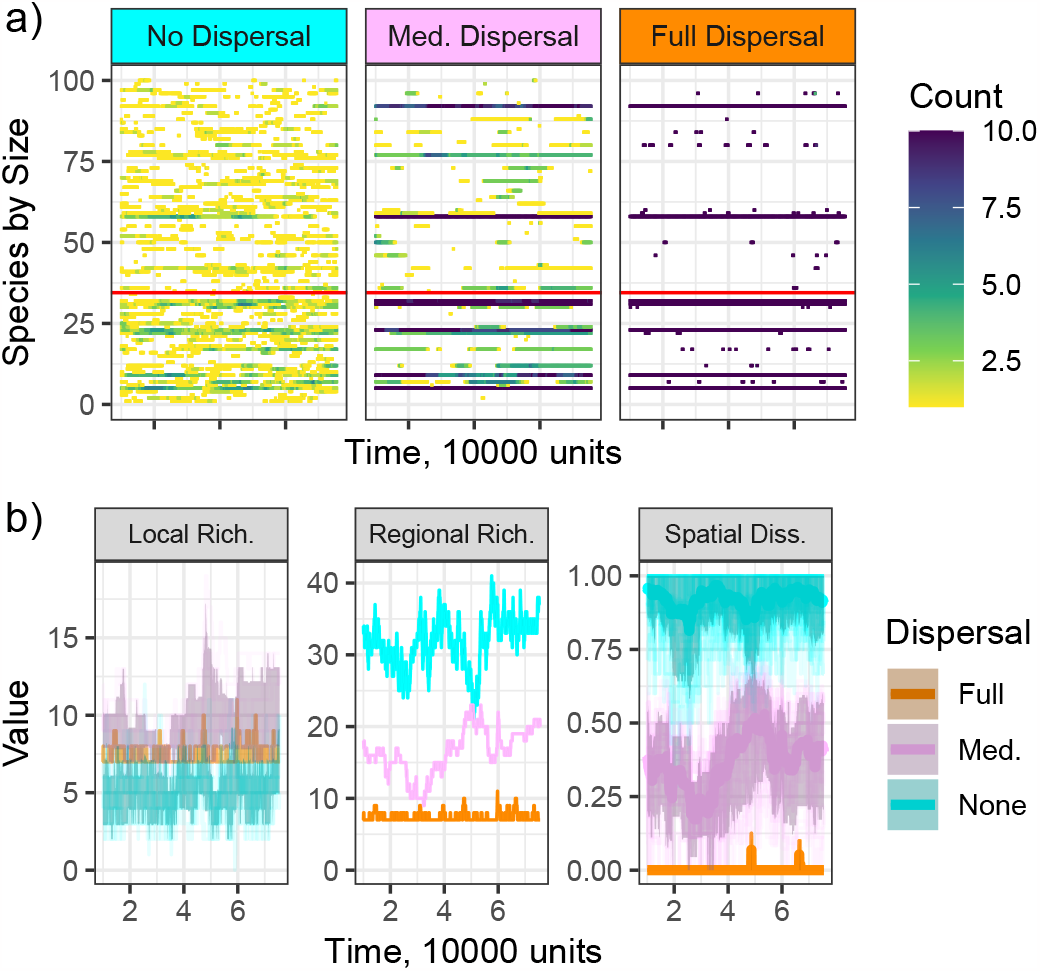
Three example simulation runs. Patch outcomes differ greatly dependent on the amount of dispersal permitted between the heterogeneous patches. Each simulation had the same neutral immigration and neutral extirpation history of 9370 events, and the same set of patch dynamics, but different dispersal rates (burnt-in for the first 10,000 time units prior to extracting results). Panels (a) show the occurrence of species, with horizontal lines indicating species identities, the duration of their presences in the patch network, and the number of patches they were present in (‘count’ colour shading) for the three dispersal rates (no dispersal: 0 dispersal rate, med.: 2 × 10^−5^, full: 0.86). Species are ranked by size, with the horizontal red lines separating basal species (below) and consumer species (above the red lines). Panels (b) show the resulting number of species per patch through time (local richness), the total number of species in the network (regional richness) and the extent to which species were shared among patches (Jaccard dissimilarity index value near zero) or communities contained different species in each patch at a given time (Jaccard dissimilarity values near 1). The dark regions in the local richness (among 10 individual patches) and spatial dissimilarity plots (in pairwise comparisons between patches) correspond to 80% intervals, whereas there is only a single value for the total number of species present in the entire metacommunity at a given time (lower middle panel).

In these three example simulations, exactly the same species immigrated at exactly the same time. This means that all of the species that successfully established in at least one patch in the zero (between patch) dispersal scenario also arrived in the corresponding patch in the full dispersal simulation, but many of them failed to establish at all. The long purple horizontal lines in Figure 3 (a) indicate communities with limited (temporal) turnover – either extinction or establishment of new species. The high dispersal simulation generates communities that are robust to further invasion and successful across heterogeneous environments (once established), whereas immigrants are more likely to establish in isolated (no dispersal) communities. When patches are homogeneous, even medium levels of dispersal are sufficient to generate high community stability and uniformity (i.e., they look like the high dispersal scenario in heterogeneous environments), shown in Figure 4. Furthermore, the time series of events (colonisations, extinctions, extirpations) at both regional and local scales show clear evidence of temporal clustering, potentially indicative of cascades of change, and this clustering increases as dispersal does (see Supplemental Information Figure S9). However, the increase in clustering with dispersal is primarily generated by some species establishing temporarily in otherwise robust communities, uninterrupted by other species (establishments followed by extinction of the same species; Figure 3 (a)). Hence, these are not ‘true’ cascades in which establishment would lead to increased rates of establishment and extirpation by *other* species thereafter.

**Fig. 4.**
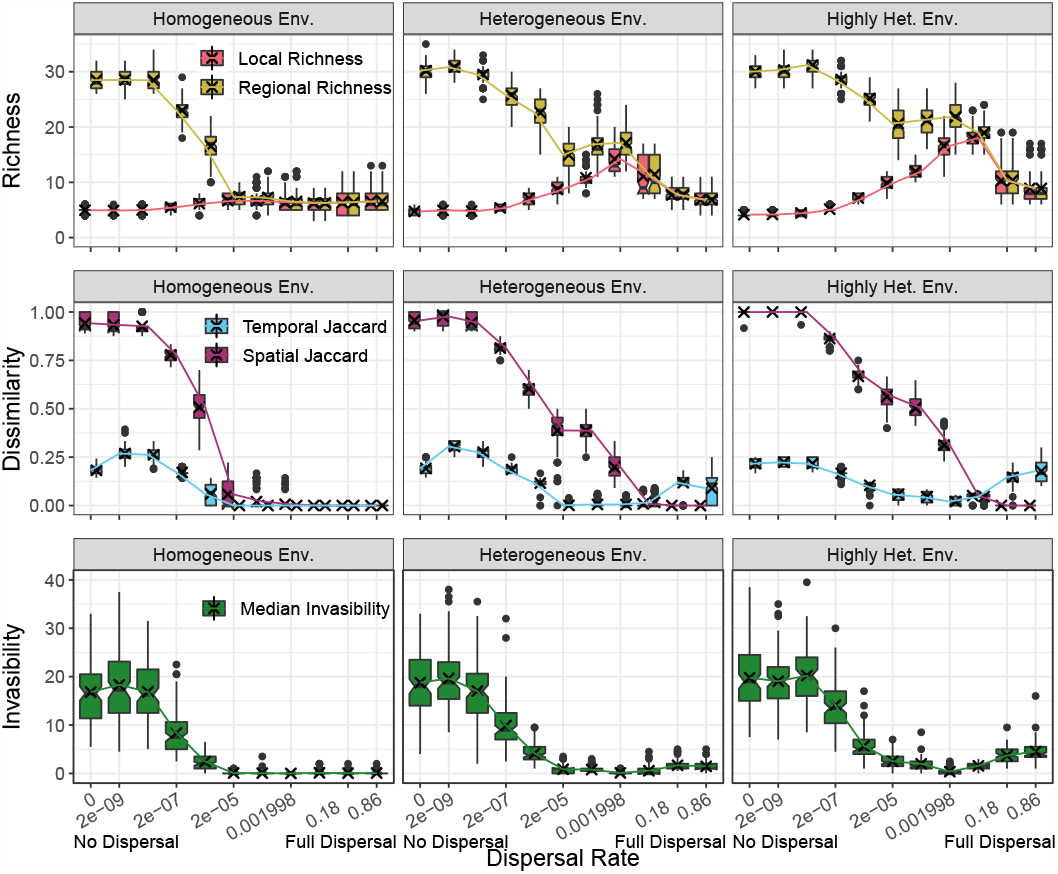
Metasimulation results for different biodiversity metrics. The species richness (top row), community dissimilarity (middle row), and invasibility (bottom row) of communities vary with the dispersal rate, but are nuanced by the amount of heterogeneity between environments (homogeneous, heterogeneous, or highly heterogeneous (left to right columns)); dispersal increases from left to right within each panel (proportions of individuals mixed between adjacent patches per time interval), covering the range from flightless sedentary organisms to true migrants. Colours indicate: local richness (top, red), regional richness (top, yellow), differences in community composition between patches (spatial Jaccard dissimilarity, magenta), how much community composition changes over time within patches (temporal Jaccard dissimilarity, blue), and invasibility (bottom, green) of the region at the end of the simulation with respect to the species pool. Each box and whisker plot is composed of a single median estimate of the corresponding metric from each of the 3300 simulations, see Materials and Methods. Each cross represents the means of the each box-and-whisker plot, and the lines connect the crosses.

These dynamics result in low species richness on patches in the zero dispersal example, but a large regional total number of species (summed across patches) because the species present differ among patches due to environmental heterogeneity and local extirpation (high spatial dissimilarity; Figure 3). In contrast, full mixing generates higher local richness, but lower regional richness because the species are shared (low Jaccard values) among patches. While medium dispersal generates intermediate regional richness and dissimilarities among islands, it has the highest richness per island (Figure 3, bottom-left panel), supporting the hypothesis that intermediate dispersal can result in the highest levels of local diversity. Dispersal is the primary driver of trends; similar plots can be constructed for the varying immigration and extirpation rates, see Supplemental Information Figures S3 - S8.

### Metasimulation Study

To investigate the influence of dispersal (mixing rate) and patch heterogeneity on local richness and regional richness, we conducted a large family of simulations in which we varied the dispersal rate, neutral immigration rate, neutral extirpation rate, pool structure, and the presence of noise in the interaction matrices. (See Tables S1 - S3 in the Supplemental Information for details on parameter variations.)

Figure 4 represents the overall relationship across 3300 simulations between dispersal rate and our metrics of diversity, ecological differentiation (temporal turnover and spatial community dissimilarity) and the resistance of the ‘end of simulation’ communities to subsequent invasion from the mainland species pool. (Additional simulation results are shown in the supplemental information, Figures S11, S12.) The relationship described in the three examples holds in general. Local richness (Figure 4 top row, red; the number of species per patch) peaks at intermediate dispersal rates (although barely when patch environments are homogeneous), confirming this hypothesis, while regional richness appears to reduce with increasing dispersal rates (top row, yellow). Broadly then, local and regional richness show opposing trends with increasing dispersal, until one reaches very high dispersal, when both local and regional richness values decline. The latter likely arises if species suited to particular patches (which differ somewhat in environmental conditions in the heterogeneous and highly heterogeneous cases) may be swamped out in runs with full dispersal. These results are robust to variations in pool parameters and immigration and extirpation rates (Figures S11, S12).

Measures of beta diversity (Jaccard dissimilarity) over both space (magenta) and time (blue) are similarly sensitive to dispersal rates (Figure 4, middle row). Spatial Jaccard falls as the dispersal rate increases, indicating that high dispersal homogenises communities, as might be expected, and seen in the three example runs (Figure 3).

Temporal Jaccard indicates the extent to which the species composition on patches changes through time. This reveals relatively high turnover in the lowest dispersal systems, indicating a greater capacity of immigrants (from the mainland pool) to establish on otherwise unconnected (isolated) patches, suggesting that dispersal among patches increases resilience. Temporal turnover reaches a minimum at intermediate dispersal rates (approximately where local richness peaks) before increasing just as local richness is falling (unless patch environments are homogeneous, Figure 4, left column). This arises in the simulations because new immigrants arrive in a single patch and quickly disperse to establish in the best (due to environmental variation among patches) patch for them, only to continue dispersing away more quickly than they can grow in abundance. The effect is related to source-sink dynamics and harvesting: population sources cannot persist if survivors cannot sustain the drain of lost (emigrant, harvested) individuals. It is only seen clearly when a substantial fraction of the total abundance of each species is redistributed among patches per time step, so this will be rare even in human-modified communities.

At the end of the simulations, we ‘challenged’ each patch-community with species that exist in the mainland pool but that were not present in the archipelago to evaluate their resistance to further ‘invasion’ (47–49). This revealed that low (among-patch) dispersal systems could be invaded much more easily than high dispersal systems. Taken together with the temporal Jaccard pattern of change, these results indicate that community networks become increasingly robust to subsequent invasion with increasing dispersal; more resilient (and commonly but not always more locally diverse) food webs are constructed from a larger pool of possible members. Invasibility and temporal turnover only diverge at the very highest (and unlikely) dispersal levels, where source-sink dynamics between heterogeneous environments as well as species pool comes into play. Thus, the low invasibility and temporal turnover experienced by high dispersal systems makes it difficult to move them out of their equilibrium state.

These overall conclusions remain consistent across a range of different assumptions including varying the immigration rate, stochastic extirpation rate, removal of stochastic extirpations, and the lower and upper limits of species sizes (Figures S3 - S8, S11, S12). Additionally, while our computational implementation of extirpation involves the random removal of a local population, the results are robust to changes to this assumption. For example, simulations where extirpation removes only 90% of a population show the same trends in the relationship between richness and dispersal (not shown). Varying dispersal consistently generates the overall shapes of the curves in Figure 4. Invasibility also increases with stochastic extinction, especially in low dispersal metacommunities. These communities lack particular components within their food webs, which in turn allows invasion. Since real island communities often lack particular functional components (either because of failed colonisation or past extinction) and are susceptible to invasion, the increase in invasibility with reduced dispersal appears realistic.

## Discussion

Our metacommunity model results indicate that increasing the rate of dispersal between ecological communities (patches, islands) produces trade-offs between diversity metrics. Local diversity (the number of species per patch, or alpha diversity) typically peaks at intermediate dispersal, while increasing dispersal drives declines in the distinctiveness of ecological communities (Jaccard dissimilarity, beta diversity). Consequently, the total number of species in the system (regional or total diversity, gamma diversity) mainly declines with increasing dispersal, although the intensity of the peak and fall depend on the amount of heterogeneity between environments. Since environments are never completely homogeneous among patches or islands, we expect the intermediate dispersal peak to be present in most if not all empirical systems. Colonisation and extirpation events are temporally-clustered (community cascades as food webs reorganise), but communities experiencing higher dispersal rates became robust to further invasion. Hence, increases and decreases in just one process – dispersal – generates a variety of community dynamics and changes in measures of diversity. There is no universally ‘best’ level of dispersal or diversity metric (50), nor do they behave in the same way as one another, as might be wished in the context of biodiversity indicators and global biodiversity policy development (9).

These results provide an opportunity to address the apparent disagreement between studies highlighting declines in biodiversity and studies highlighting variable but no-net decline trends. Although humans have altered ecological communities in many different ways, altered rates of dispersal are a common consequence. This frequently results in a more well-mixed world. The establishment of ‘non-native’ species is accelerating globally (51). Equally, ‘native’ species are increasingly moving: range boundaries generally are shifting polewards and to higher elevations (and deeper depths) in response to anthropogenic climate change (e.g., 52–54), such that species are colonising ecological communities from which they were previously absent at unprecedented rates. In contrast, metapopulation ecology has shown that dispersal (colonisation) rates of habitat-specific species declines with habitat isolation, e.g., from habitat fragmentation (55), although the establishment rates of edge- and disturbance-related species may increase (56).

This means that the expected outcome in recent biodiversity monitoring is context-dependent. It depends on the measure(s) of diversity change (as above), how isolated a community was prior to human intervention (isolated island communities have experienced most extinctions associated with ‘non-native’ species), whether dispersal rates have increased or decreased in relation to local environmental changes, how different the local environments are, and the types of species considered. For example, habitat core and edge species may experience opposite changes to their dispersal rates associated with their abilities to move through human-modified environments (many studies focus on core species). Species have different likelihoods of being transported by humans, and the natural dispersal rates of different taxonomic groups vary over many orders of magnitude (many studies are of only one or a few groups). Hence, different taxa may show different diversity trends even for identical sites. A combination of these factors should contribute to the debate (see the Introduction) over exactly what biodiversity trends exists.

Our results provide similar context–dependent insights into the ‘invasion debate’ (57–60). The model simulations exhibit emergent series of temporally correlated events (establishments, extirpations; though often arrivals followed by disappearance of the same species) consistent with narratives of community cascades (or meltdown; 61) and the extinction of endemic species when previously-isolated communities experience increased immigration (e.g., 13, 62). Nearly all IUCN-listed threatened species that are endangered by invasive (i.e., not previously present) species are in previously-isolated communities: island communities for reptiles, birds, mammals and plants; and ‘island-like’ wetland environments within continents for freshwater fish and amphibians (63). In contrast, many other ‘natural’ ecosystems in continental regions (which have had higher background immigration rates) appear relatively robust to invasion, compared to human-altered ecosystems (which may still be in the ‘burn-in period’) (e.g., 64, 65).

This simulation study has proposed one simple mechanism, dispersal relative to spatial scale, that has the capacity to connect the literature on the decline of global biodiversity, on how local biodiversity can remain relatively stable on average, and on how isolated ecological communities experience reorganisation (including extinctions but also increased community-level diversity) when experiencing recently-increased immigration rates. While we find that dispersal is sufficient to explain different sides of the diversity and invasion debates, dispersal also interacts with many other temporal changes to the environment and these pressures should be considered together in any discussion of policy options. Whether people wish to manipulate or control dispersal rates will depend on which components of diversity and community invasibility, as well as functional processes, are prioritised in a given context.

## Materials and Methods

Pseudocode for the core processes is provided in the supplemental information for transparency and replication (66), and the full simulation code is available (see Data Availability).

In what follows, we describe the construction of our model mathematically, beginning with the (singular) pool and average interaction matrix. We then discuss how we combine multiple patches, dispersal, and the inclusion of stochastic events. We describe the numerical evaluation of our model, before finishing with a discussion of how we varied parameters and analysed results.

### Pool and Interaction Matrix Construction

A mainland (species pool) - island (patch, local community) framework was selected to consider the impacts of different (inter-patch) dispersal rates on the balance between local community diversity (patch-level *α*) and regional (all patches *γ*) scales, and the compositional differences between communities (*β*). This approach was both computationally tractable and relevant to the real world, where numbers of species per oceanic island have often grown with increased human-assisted dispersal (increased *α*), while the extinction of endemic species (reducing global *γ*) associated with biological invasions (new community interactions and structures) are greatest in such locations (13, 67). The methodology for constructing a species pool and the notation (including the numbering on parameters *k*_1_ through *k*_6_) used follow previous literature by Law and Morton (37), whose approach was informed by Cohen et al. (68).

A pool of 100 species, 34 basal and 66 consumer, is constructed by first assigning each species a number *q*_*i*_ drawn from uniform distributions on (−2, −1) if basal and (−1, 0) if a consumer unless otherwise specified. The size of species *i* is then *s*_*i*_ = 10_*q*_*i* (hence-forth labelled grams, but in principle representing a general measure of size (37)). Basal species have positive intrinsic growth rates and negative intraspecific interactions, but do not interact with other basal species. As a result, basal species abundance decreases primarily as a result of consumer consumption. Consumer species eat anything smaller than themselves (but have a built-in size preference as in real world food webs (68)), have negative intrinsic growth rates (that result in death if they run out of prey), and have no intraspecific interactions. Both basal and consumer species can be eliminated via too much consumption by their own consumers.

In what follows, we provide an explicit mathematical description of how the average growth rate (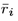 for species *i*) and interaction terms (*ā*_*i,j*_ for species *i* and *j*) are calculated. We then explain how these terms can be modified between patches to allow patch-level environmental heterogeneity to be captured.

### Intrinsic rate of increase

To construct species *i*’s intrinsic growth (or loss) rate *r*_*i*_ (units per day), the mean is set to

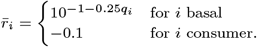

The basal intrinsic growth rate is empirically motivated (69, 70): a linear relationship is fitted to log *r* as a function of log body weight (see appendix of 37). Consumers rely on consuming prey to balance their intrinsic loss rate in order to survive/avoid starvation.

### Species intra-actions and basal biomass

To construct basal species *i*’s intraspecific interactions (*a*_*i,i*_), we set the mean to

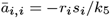

where *k*_5_ = 100 (grams times species abundance density) determines the basal equilibrium biomass. This is so that each basal species has the same maximum carrying capacity when measured in terms of biomass with no consumers and we thus constrain the maximum basal biomass that a patch supports to 3400 grams per volume (patch). As noted above, consumer species *i* has *a*_*i,i*_ = 0 (i.e. no intra-specific competition or density dependent growth).

### Species interactions

To construct average species interactions *ā*_*i,j*_, *i* ≠ *j*, we compare the relative sizes and types of the species. We take the constants *k*_1_ = 0.01 (units per time and species abundance density) to be the scale of the interaction strength relative to the growth rates, take *k*_2_ = 10 to be the preferred size ratio of predator to prey, take *k*_3_ = 0.5 to be the willingness to deviate from the preferred prey size, and take *k*_4_ = 0.2 to be the energy efficiency from consuming prey (37).

Then the mean effect of species *j* on species *i* is

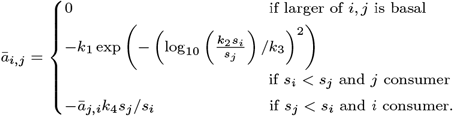

Here, a basal species does not interact with species smaller than itself. If the larger species *j* is a consumer, then the inner expression of 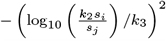 provides diminishing returns as the prey deviates from the consumer’s preferred size: *k*_2_ determines the location of the maximum and *k*_3_ the penalty for deviation. The size ratio and efficiency *k*_4_ are then used to convert the abundance density from prey species to consumer species. Note that the gain to a consumer from consuming the prey and the loss to the prey from being consumed are not independent and are of opposite sign.

### Lotka-Volterra Model

The average interactions *ā*_*i,j*_ then can be used as a 100 × 100 Lotka-Volterra interaction matrix. In this case, the model in the absence of dispersal and environmental heterogeneity is

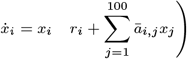

for species 1 ≤ *i* ≤ 100 with the abundance density of species *i* written as *x*_*i*_ and its time derivative as 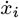. The term 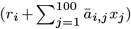 is the per capita growth rate of species *i*, which can be used to determine if species *i* could grow from a small population. Finally, we note that this system of equations captures all community- or patch-scale ecological dynamics with linear responses.

### Parallel Community Assembly

We now highlight our additions to and points of departure from the Law and Morton model of the previous section.

### Multiple patches

Instead of considering only a single community assembly process, we now construct multiple communities which model small environmental differences between discrete patches. We set a number of patches, here 10, fix the species pool, and then used the mean interactions *ā*_*i,j*_ exactly as defined above, to generate patch specific interactions _*p*_*a*_*i,j*_ (unless there is no environmental heterogeneity).

For a patch *p*, each element of the associated interaction matrix is given by a random sample from a normal distribution truncated at 0 to preserve the sign of the element. That is

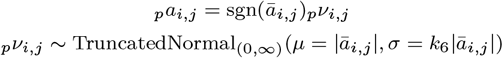

where sgn is the sign function, TruncatedNormal_(0,∞)_ is the truncated normal distribution (i.e. the distribution conditional on the event that a draw is from the range (0, ∞) (71), implemented in the `rtnorm R` package (72)), | · | takes the absolute value, and *k*_6_ = 0.1 is the coefficient of variation of the underlying normal distribution. Since *k*_6_ = 0.1, we expect all interaction matrices to be broadly similar, but to differ in exact magnitude (but not the sign of the interaction). Our highly heterogeneous environments case doubles *k*_6_ to 0.2, while the homogeneous case takes _*p*_*a*_*i,j*_ = *ā*_*i,j*_. (All draws from the truncated normal distributions are independent.)

As all patches in a given run of the simulation share the same pool, we need not vary the pool in the same way. We still allow for variation in the parameters of the basal and consumer species by sampling the intrinsic growth/loss rates analogously with a fixed coefficient of variation:

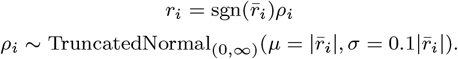

### Dispersal between patches

We also allow movement between patches following a network that is, in principle, arbitrary but in practice set to a one dimensional torus (a ring). For each pair of patches we set a (positive) resistance to dispersal defined between them (taken, but not required, to be symmetric). If the resistance is infinite, no dispersal can occur directly between those two patches. We take the lower bound in our simulations to be a resistance of 1, at which ‘full dispersal’ takes place. Each species is then assigned a ‘speed’ (taken to be 1 for all species here), and together these define a dispersal (rate) matrix *D* which describes how quickly species abundance densities move between patches. We take the diagonal of *D* to be the negative of the (non-diagonal elements of the) column sums to conserve mass. Note that *D* is banded, corresponding to preventing abundance from changing species when dispersing from one patch to another. Hence, for species *i* on patch *p*, the rate of change of its abundance density is

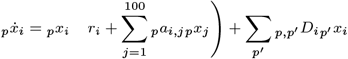

where the additional labels capture the patch and the additional term captures movement of species *i* to and from patch *p*.

### Events and time scales

The Law and Morton model results in final, static, uninvasible communities (or cycles of communities) using time scale separation and by alternating between community dynamics and migration dynamics. We instead simultaneously assemble multiple interconnected communities to examine how they change through time. In addition to (neutral, local) immigration events, we first add their complement: extirpation (neutral, local extinction) events. This replicates local stochastic disasters. The implementation is similar for both. For immigration events, a species *i* is selected uniformly at random from the pool and a destination patch *p* is selected uniformly at random from the patches. If the per capita growth rate (see Lotka-Volterra Model, above) is positive a small initial population is added to that patch. (The size of the initial population is discussed below, see Solver abundance details.) For extirpation events, if the species *i* is present on patch *p*, it is instead removed by setting its local abundance density _*p*_*x*_*i*_ to 0.

We numerically resolve the differences in time scale between immigration and extirpation events and community dynamics, we assign to each event an exponential waiting time. The exponential distribution’s rate is the magnitude of the largest eigenvalue of the interaction matrices multiplied by a multiplier (taken to be 0.1, 1, or 10 independently for each as a simulation parameter, as seen in supplemental Figure S11). The largest eigenvalue of the interaction matrices can be interpreted as the natural rate at which the interactions as a whole take place. Hence, the events should take place on a similar time scale as the fluctuations induced by community dynamics.

### Numerical Evaluation

Unlike Law and Morton, we are numerically resolving the community dynamics on the same time scales as immigration and extirpation, so we do not use permanence methods requiring time scale separation (alternating resolving a migration or extirpation event and allowing dynamics to proceed to a community equilibrium (see 37, 73, 74)). Instead, we numerically integrate the system using R’s deSolve package, which implements ODE-PACK functionality including the capability of mixing the evaluation of events with the evaluation of a dynamical system (75–77). We primarily use the lsoda algorithm, which allows for events and can alternate between stiff and non-stiff methods for numerical integration (78).

### Solver abundance details

In order to use this solver, we must specify how the continuous (time, abundance) interacts with the discrete (immigration, extirpation). Relating abundance to our events first, we specify that at the times when abundance densities are reported, we remove species *i* on patch *p* if its abundance _*p*_*x*_*i*_ *< E* = 10^−4^ by setting _*p*_*x*_*i*_ = 0. This allows for community dynamics to induce extirpation, while avoiding problems of numerical precision. This does, however, implicitly specify an “individual” scale for our simulations: _*p*_*x*_*i*_ = 1 corresponds to 1*/E* = 10, 000 individuals. As discussed above, we also set species *i* in patch *p* to 0 if a neutral extirpation event occurs. On the other hand, if a neutral immigration event occurs and the species would have positive per capita (local) growth, colonisation occurs and species *i* is added with initial abundance density 4 × 10^3^*E* = 0.4. This is approximately in line with the minimum viable population size (79).

### Solver time step details

In order to reliably detect community dynamics driven extirpations, we check for extirpations between events over either ten equal sized intervals spanning the waiting time between events or every unit (length 1) time interval if the waiting time is longer than 10 time units. Frequent checks between events for community driven extirpation also forces the algorithm to use more steps, resulting in smoother results. The trade-offs are increases in the time required to finish the simulation (due to frequent interruptions), increases in the file size (more entries are reported), and the additional time steps are unnecessary if community driven extirpation does not occur (time steps are otherwise dynamically and internally calculated by the solver). Additionally, we burn-in the simulations used in the metasimulation study, discarding times before 1 × 10^4^ time units and uniformly end the simulations at 6.5 × 10^4^ time units. The former allows the system to approach equilibrium, while the latter prevents large disparities in simulation length due to random fluctuations in event sampling.

### Meta-analysis

Our results in Figure 4 include only some of the parameter variations simulated. Here, we detail how the various runs differ from each other. Parameters and their combinations are given explicitly in Tables S1 - S3. We also note our usage of `R’s tidyverse` package for data handling and plotting (80), the `patchwork` package for plotting (81), and the foreach package for parallelisation (82).

### Number and configuration of simulations

For a given parameter set, 10 different pools, each with 10 different patches, are generated. For simplicity, patches are configured in a ring (i.e. *↔* 10 *↔* 1 *↔* 2 *↔* …) whose links are each assigned a single dispersal value; see the construction of *D* above. Each pool is then assigned 10 independent histories, which define 10 different simulation runs, allowing for the possibility of exploring pool-related emergent structure. Each run then contains 10 *α* values through time, 1 *γ* value through time, 45 spatial *β* values through time, 10 temporal *β* values through time (we fix the time difference to 1*/*100th the simulation length for computational efficiency), and 10 patch-level end of simulation invasibility calculations. These are all derived from the presence and absence of species in the 10 communities in the system.

### Variation of parameters

We vary the following parameters: the basal size lower bound, the consumer size upper bound, the coefficient of variation when sampling the interaction matrices, the (multiplier on the) rate of immigration events, the (multiplier on the) rate of extirpation events up to the exclusion of such events, and the rate of dispersal (see Table S2). All other parameters in the system are fixed (see Table S1). We provide a full table of the parameters and how we vary them in combination in the Supplemental Information (Table S3). We vary these parameters as a robustness check on our parameter choices and conclusions. As discussed in the main text, we provide alternative plots in the Supplemental Information. We finish with *∼* 27, 900 simulations completed. The remainder (*∼* 7, 500) failed due to memory or computing time constraints as discussed in the Supplemental Information.

To combine the results into Figures 4, S10, and S11, for each simulation we take the median values of *α*, spatial *β*, and temporal *β* (amongst all patches and over all times between 1 × 10^4^ and 6.5 × 10^4^ time units) and median values of *γ* (over all times) and end of simulation invasibility (see below). Figure 4 is then composed of these values for simulations with immigration and extirpation rate multipliers of 1 and no changes to the pool parameters (see Pool and Interaction Matrix Construction), with values aggregated by dispersal and interaction matrix coefficient of variation and presented using standard notched box-and-whisker plots (83) and means (shown as lines and crosses). This prevents simulation length from biasing the analysis.

### Jaccard Index

In order to quantify how patches differ from each other within a time step (spatial dissimilarity, spatial *β*) or how patches differ from themselves across time (temporal turnover, temporal *β*), we use the Jaccard (dissimilarity) index on the corresponding presence absence data from our simulations as implemented in the vegan R package (84, 85). Specifically, if *A* is the set of species present in one patch and *B* is the set of species present in a different patch at the same time (spatial *β*) or the same patch at a different time (temporal *β*), then we calculate

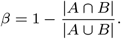

### Quantifying Invasibility

Quantifying *α* and *γ* is straightforward; we identify which species are present within patches and within the overall system. Quantifying invasibility is less clear. We quantify invasibility as the number of species in the pool that could both establish in a patch (positive local per capita growth rate) and then subsequently survive the full community and dispersal dynamics for 5 times the inverse of the magnitude of the largest eigenvalue of the interaction matrix (which is the inverse of the rate of interactions and events and thus sets a natural time scale). The invasibility values reported in Figure 4 then correspond to the median invasibility of a patch (computed over all patches) at the end of the simulation.

## Data Availability

All code, random seeds, R scripts, and summarised data used to generate the simulations and subsequent figures are available at https://github.com/Brennen-Fagan/Community-Assembly. The > 10 TB of data generated from the code, random seeds, and R scripts are available by request from the corresponding author.

## ACKNOWLEDGMENTS

This work was funded by a Leverhulme Trust Research Centre - the Leverhulme Centre for Anthropocene Biodiversity. We thank the University of York Complexity and Stability Reading Group and the Sheffield Spatial Ecology Workshop, especially G. W. A. Constable, R. Law, T. Carroll, W. F. Fagan, and J. Matthiopoulos, for helpful discussions. We also thank M. Vellend, C. Dytham, and J. Hatfield for valuable comments on the article.

The majority of this project was undertaken on the Viking Cluster, a high performance compute facility provided by the University of York. We are grateful for computational support from the University of York High Performance Computing service, Viking and the Research Computing team.

