## Supplemental Information for "Increased dispersal explains increasing local diversity with global biodiversity declines"

#### Increased dispersal can explain the paradox of increasing local diversity while global biodiversity declines

##### This PDF file includes:

Supporting text
Figs. S1 to S13
Tables S1 to S3
SI References

#### Supporting Information Text

### Contents


**1 Convergence of Simulations** **3**

**2 Example Metrics over Time** **3**

**3 Local and Regional Richness Comparison** **3**

**4 Variation of Neutral Rates, the No Extirpation Case, and Variation of Assembly Parameters** **4**

**5 Pseudocode** **5**

A Notation . . . . . 5

A.1 Basic types . . . . . 5

A.2 Structured types . . . . . 5

A.3 Basic functions . . . . . 5

B A typical run and actual use . . . . . 6

C Algorithms . . . . . 7

**6 References, Tables, and Images** **13**

S1 System parameters that are varied between simulations. . . . . . 13

S2 System parameters that are varied between simulations. . . . . . 13

S3 Sets of Simulations Conducted. . . . . . 14

S1 Simulations Completed - Absolute . . . . . 15

S2 Simulations Completed - Relative . . . . . 16

S3 Presence-Absence, No Dispersal . . . . . 17

S4 Presence-Absence, Med. Dispersal . . . . . 18

S5 Presence-Absence, Full Dispersal . . . . . 19

S6 Presence-Absence, Local Richness . . . . . 20

S7 Presence-Absence, Regional Richness . . . . . 21

S8 Presence-Absence, Jaccard Distance . . . . . 22

S9 Inter-event Time Interval Length . . . . . 23

S10 $\alpha$ - $\gamma$ state space . . . . . 24

S11 Metrics by Neutral Rates . . . . . 25

S12 Metrics by Pool and Noise Parameters . . . . . 26

S13 Diagram of an Example Simulation Run. . . . . . 27

#### 1. Convergence of Simulations

Of the 35,400 simulations that we aimed to conduct, 27,912 were able to complete due to memory and time constraints. Particular parameters, their values, and their combinations can be seen in Table S1, Table S2, and Table S3. In general, more events meant that the simulations were slower, leading to discrepancies in which systems were most fully explored. We can characterise this with heat maps, see Figure S1 and Figure S2 for absolute and relative comparisons of the simulations completed. Most of the missing simulations are indeed when at least one of (the) extirpation or immigration (rate multipliers) is high. This is complicated by additional computational complexity in the low dispersal rate cases (where dispersal changes underlying communities but is not sufficient to homogenise them).

With that having been said, this does not substantively affect our analysis. While lower percentages of simulations are completed with high numbers of events or low amounts of dispersal, our simulations were designed with good coverage of the ratio between immigration and extirpation rates and we show in Sections 2 and 4 that the results remain generally consistent for different neutral rate multipliers.

#### 2. Example Metrics over Time

In Figure 3 of the main text, we showcased how dispersal affects three example simulations with heterogeneous environments, with the same extirpation and immigration histories, and with both extirpation and immigration set to the same rate. In Figure S3 - Figure S8, we reproduce Figure 3 for each combination of immigration and extirpation rates to allow for visual comparison of simulations. Note that changing the rates necessarily changes the history of events, so simulations with the same immigration and extirpation rates will have the same history (i.e., between Figure S3, Figure S4, and Figure S5), but simulations with the same dispersal rates will not (i.e., within Figure S3, Figure S4, or Figure S5).

Figure S3 - Figure S5 show a few differences that are worth commenting upon. First, higher immigration and extirpation rates are necessary to maximise the coverage (minimise the white space) of the presence-absence plots. Second, higher extirpation and lower immigration rates do inhibit the formation of a core community. Third, changes in either rate on its own changes the species that are observed, possibly due to perturbing the system into a different core community. Despite these differences, generally the same communities appear to be forming indicating that these are mostly controlled by the pool and dispersal rates.

Similarly, the behaviours in Figure S6 - Figure S8 show us that the story within each panel is consistent between each panel, even as the variations between each panel are similar. No dispersal results in communities with consistently less local richness than full dispersal systems, which are generally comparable to systems with medium dispersal. Regional richness reverses this, with no dispersal coming in above medium or full dispersal except for when immigration is far outpaced by extirpation. Finally, dissimilarity between patches within a system is maximised when there is no dispersal and minimised when there is full dispersal.

We close this section by commenting on temporal clustering in Figure 3. To assess the possible temporal clustering associated with cascades of events (here including events due to stochastic immigration or extirpation, community dynamics, or inter-patch dispersal), the following time series analysis was performed: for each time series of events shown in Figure 3 (a), lists of event times were extracted at the scales of island ("Local", all successful events regardless of cause), archipelago ("Regional", i.e., successful immigration to or extirpation from the entire region, regardless of cause), and mainland ("Neutral", all stochastic neutral events regardless of success). These events were then double-checked against the list of neutral events to categorise them. The inter-event time intervals were then analysed to ascertain whether non-clustered (exponential) or clustered (gamma) distributions better fit the observations. Fitting was performed by maximum likelihood estimation in MATLAB after adding noise (smaller than the smallest time interval) to 0's (resulting from discretisation of recording times) (1). The results are in Figure S9. As anticipated, the neutral (pool) events are identical (representing the same history) and are distributed exponentially (Figure S9, first row). For the no dispersal cases (Regional, No Dispersal and Local, No Dispersal) the time series of events is again approximately exponential (Figure S9, first column). There is a slight trend towards temporal clustering gamma with medium dispersal (Figure S9, second column) and clear evidence of temporal clustering in the full dispersal cases (Figure S9, third column).

#### 3. Local and Regional Richness Comparison

Dispersal's effect on our simulations operates on a continuum, Figure S10 (base case in the sense of Table S3 with coefficient of variation set to 0.1). At very high dispersal rates (rightmost column), the core community emerges rapidly with nearly no variation, reflected in a very tight linear relationship between regional richness and local richness. Each patch is essentially equivalent in terms of species presence (low Jaccard dissimilarity). As the dispersal rate decreases (moving leftward), we see more variation in the patches. There is still a physical boundary as regional richness must be greater than or equal to local richness and both must be non-negative, but the individual local and regional values from our simulations spread out along and slightly away from this boundary, although the exact manner of spreading depends on the other parameter values in non-trivial ways. This behaviour continues with the departure from the physical boundary increasing as dispersal continues to decrease until the system no longer touches the physical boundary with high probability. At very low dispersal rates, the system is mostly subject to the effects of the neutral immigration and extirpation events and the resultant community dynamics.

#### 4. Variation of Neutral Rates, the No Extirpation Case, and Variation of Assembly Parameters

The original community assembly model of Law and Morton (2, 3) does not include any of parallel assembly and a single time scale, dispersal between parallel communities, or neutral extirpation events. The problem of time scales can be investigated by making events sufficiently rare and with no dispersal. In this case, the system stabilises faster than events arrive in the system and recreate the (local richness) results of Law and Morton's original work (2). Additionally, our focus in the main text is on dispersal. In this section, we discuss the effects of neutral extirpation events as well as the influence of the original size scale and noise on the various diversity metrics.

To do so, we begin by considering variations of Figure 4 generated using different combinations of neutral parameters in Figure S11. As in Figure 4, Figure S11 displays the relationships between local and regional measures of species richness (top rows), temporal and spatial measurements of Jaccard dissimilarity (middle rows), and the invasibility of the system with respect to the pool at the end of the simulation (bottom rows). Additionally, we break each of these up by the extirpation (Ext., columns) and immigration (Imm., rows) rate multipliers, where Ext.: 1 and Imm.: 1 is the base case shown previously in Figure 4 (middle column).

The trends are generally similar between panels, even within panels missing simulations (see Section 1). Local richness (top, red) rises until dispersal rates of about 0.002, after which it begins to fall. Regional richness (top, yellow) generally falls as dispersal rates rise. Similarly, spatial Jaccard dissimilarities, measuring the between patch inhomogeneity at a given point in time, falls as dispersal rates rise. Temporal Jaccard dissimilarities and end of simulation invasibility, on the the other hand, generally fall or remain low for small dispersal rates. For our highest dispersal rates, they instead rise again, as discussed in the main text. This rise is smaller for invasibility and the temporal Jaccard dissimilarity is partially controlled by the amount of immigration. Figure S11 makes clear that the no extirpation case lies on a natural continuum that can be explored more fully by considering the full range of neutral dynamics rather than just neutral immigration.

In order to examine the effects of the pool's size range, meaning how large we allow the largest consumers to be and how small the smallest basal species, and the effects of noise, referring to whether we randomly draw each patch's interaction matrix elements from normal distributions or instead take the mean value for each, we again break apart Figure 4. This time, we break up the panels of Figure 4 by the pool and noise combination, see Figure S12. The combinations of pool and noise, from left to right, refer to the base case, highly heterogeneous environments case (i.e., doubled coefficient of variation of the interaction matrices), homogeneous environments (i.e., no noise in the interaction matrix), to larger consumer species (i.e., an increase of 1 to the maximum logarithmic body size), to larger consumer species and smaller basal species (i.e., a decrease of 1 to the minimum logarithmic body size), and to smaller basal species. Each size change is by one on the logarithmic scale, and thus one order of magnitude on the actual size scale.

There are some modest observations of note in Figure S12. Proceeding in the usual fashion, local richness as a function of dispersal seems to have the same trend of rising and then falling as it reaches agreement with regional richness, but there is a small amount of variability stemming from the pool or noise. For low amounts of dispersal, the base and large consumers cases have mild decreases in local richness compared to the cases that allow for smaller basal species. Such smaller basal species would be less easily predated by consumers, which might explain the higher local richness, but does not immediately explain the pattern in regional richness, where the base case generally has the most richness and the larger consumers and smaller basal species case has the least. We conjecture that the base case has the best balance of basal and consumer species. Since the smaller basal species are less easily predated, the consumers are less able to receive food (since the number of basal species and total basal biomass are fixed, meaning some of the basal biomass is less accessible due to size differences). While the same issue exists for larger consumers, larger consumers would instead rely on consuming other consumers (due to the default choice of food size preference). Meanwhile the homogeneous patch environments case uniquely experiences a precipitous decline in regional richness with an almost invisible increase in local richness, as there are no differences in patches to facilitate species diversity. Finally at maximum dispersal, there is very little distinction between the different pool configurations and noise structure for richness.

Continuing to spatial Jaccard, we observe the comparably sudden decline in the homogeneous patch environments case matching its decline in regional richness as well. Reflective of $\beta$ diversity's relationships with $\alpha$ and $\gamma$ diversities, spatial Jaccard is consistent with our observations above regarding local and regional richness with smaller basal species compared to larger consumer species. Smaller basal species, and less accessibility for consumer species, then drives the comparatively smaller values for temporal Jaccard via rapid assembly and inability to place more consumers that might overturn the system. Turning to the high dispersal set of systems, we see resurgence primarily in the base, larger consumer species, and smaller basal species cases. In contrast, the smaller basal and larger consumer species case still has very little in terms of species that can colonise successfully, while the homogeneous patch environments case has effectively none. This suggests that the inhomogeneity between patches and the rate of dispersal are driving the temporal turnover; species arrive on a patch, quickly find the best patch, but then are overdispersed away from their preferred environment. Similar trends can be seen in end of simulation invasibility, but the overall patterns are more similar. Despite a number of specific differences associated with variation in rates (Figure S11, Figure S12), the overall qualitative patterns for richness, dissimilarity, and invasibility seen in the main text Figure 4 holds for most areas of parameter space.

#### 5. Pseudocode

#### List of Algorithms


1 Parallel Community Assembly (PCA) . . . 8
2 Pool Construction . . . 9
3 Interaction Matrix Construction . . . 10
4 Construct per capita ecological dynamics . . . 11
5 Identify approximate characteristic rate. . . . 11
6 Construct the assembly sequence events . . . 11
7 Construct spatial dynamics . . . 12


##### A. Notation.

***A.1. Basic types.*** We use the following notation for basic objects/types:

- Nat : the natural numbers, $\{0, 1, 2, \ldots\}$
- Real : the real numbers
- Int : the integers, $\{\ldots, -2, -1, 0, 1, 2, \ldots\}$
- subscripts indicate restrictions on the sets
- DF : data frame, see below
- Func : a lambda function
- 1D-Object : any object with 1 dimension (such as a list or 1d array)
- 2D-Object : any object with 2 dimensions (such as matrices and data frames).

***A.2. Structured types.***

- A data frame (DF) is a 2D-Object whose rows represent observations or individuals and whose columns are vectors of features of each observation or individual. The values in a given column all have the same type; values in different columns may have different types. Columns may have names.
- Indexing of objects starts at 1.
- The $i$-th element of a 1D-Object $x$ is $x[i]$.
- The $i$-th row of a 2D-Object $x$ is $x[i, ]$.
- The $i$-th column of a 2D-Object $x$ is $x[, i]$. A named column can also be accessed using its name: for column '*colname*', the column is also given by $x[, \text{'}colname\text{'}]$.
- The element the $i$-th row and $j$-th column of 2D-Object $x$ is $x[i, j]$.
- Multiple elements, rows, or columns of an object can be given by replacing $i$ above with the range desired. In particular, the first $n$ elements of 1D-Object $x$ are given by $x[1 : n]$, the first $n$ rows of 2D-Object $y$ are given by $y[1 : n, ]$, and the first $n$ columns of $y$ are given by $y[, 1 : n]$.
- Addition and element-wise multiplication are vectorised. For example, if $\mathbf{a}$, $\mathbf{b}$ are 1D-Objects of the same type, structure and number of elements, and $c$, $d$ are scalars, then $\mathbf{a} \times \mathbf{b} = (a_1 b_1, a_2 b_2, \ldots)$, $\mathbf{a} + \mathbf{b} = (a_1 + b_1, a_2 + b_2, \ldots)$, and $c + \mathbf{b} \times d = (c + b_1 d, c + b_2 d, \ldots)$, each with the same structure as $\mathbf{a}$ and $\mathbf{b}$.
- For brevity, when creating an empty square matrix, we provide the number of rows as a subscript (as the number of columns is then implied). Element values default to 0.

***A.3. Basic functions.*** We use the following basic pre-defined functions:

- $\mathsf{length}$ : 1D-Object $\rightarrow$ Nat: the length (number of components) of a 1D-Objects.
- $\mathsf{rows}$ : 2D-Object $\rightarrow$ Nat : the number of rows of a 2D-Object.
- $\mathsf{cat}$ : 1D-Object $\times$ 1D-Object $\rightarrow$ 1D-Object : concatenation of 1D-Objects; for example, given $\mathbf{a}$, $\mathbf{b}$ : 1D-Object (not necessarily of the same length), then $\mathsf{cat}(\mathbf{a}, \mathbf{b}) = (a_1, a_2, \ldots, b_1, b_2, \ldots)$.
- $\mathsf{repeat}$ : 1D-Object $\times$ Nat $\rightarrow$ 1D-Object: concatenates the 1st argument a number of times equal to the 2nd argument. This preserves the original ordering in the repetitions. E.g., $\text{repeat}((1, 2, 3), 3) = (1, 2, 3, 1, 2, 3, 1, 2, 3)$.
- $\mathsf{LSODA}$ : $\text{Real}^m \times \text{Real}^n \times \text{Func} \times \text{DF} \times \text{Real}_{>0} \rightarrow \text{Real}^{(m+1) \times n}$: the call to the ODE solver, providing the starting abundance (1st argument), reporting times (2nd argument), the dynamics (3rd argument), the events (4th argument), the zeroing threshold (5th argument). This returns a matrix whose first column contains the reporting times, whose first row is the starting time and initial abundance, and whose remaining entries are the abundances through time.
- $\mathsf{rnorm0trunc}$ : $\text{Real}^n \times \text{Real}^n_{>0} \rightarrow \text{Real}^n$: sample $n$ times from a normal distribution truncated at 0, where $n$ is the length of the 1st and 2nd arguments (which should be the same). The remainder of the distribution(s) is rescaled. The 1st and 2nd arguments correspond to the mean and standard deviation of the untruncated normal distribution(s).
- $\mathsf{rpick}$ : $\text{Nat}_{>0} \times \text{Int} \times \text{Int} \rightarrow \text{Int}^{\text{Nat}_{>0}}$: sample from a discrete uniform distribution over Int a number of times equal to the 1st argument. The distribution is taken to start at the 2nd argument and end at the 3rd argument.

- runif : $\text{Nat}_{>0} \times \text{Real} \times \text{Real} \to \text{Real}^{\text{Nat}_{>0}}$: Same as rpick, but for a continuous uniform distribution.
- sortedOrder : 1D-Object $\to \text{Nat}_{>0}^{\text{length(1D-Object)}}$: returns the indices of the elements of the argument required to sort the argument. For example, sortedOrder$((3.5, 99, 1)) = (2, 3, 1)$.
- sign : Real $\to \{-1, 0, 1\}$: returns the sign of the argument, or zero if the argument is zero

**B. A typical run and actual use.** Figure S13 and the Example Run box detail the series of steps that we expect to be taken when running our code, available at https://github.com/Brennen-Fagan/Community-Assembly. The code there includes the functions mentioned in Algorithms 1 - 7 in the R folder, in addition to some functions that either support the primary functions or were used for analysis. The main functions of the project can be found in LawMorton1996SubFunctions.R and MultipleCommunityAssembly.R.

---

Example Run.

```
Choose/set Law and Morton (1996) parameters P = {Strg, Pref, Spec, Effi, EqDn, Spread}, B_min, B_max, C_min, and
    C_max (interaction strength, consumer body size preference, consumer prey nonspecificity, predation energy efficiency, basal
    equilibrium biomass density, and minimum and maximum basal and consumer body size).
Choose numbers of basal species B, consumer species C, environments N_env, immigration events N_ar, elimination events
    N_el, functions to sample immigration event times rFN_ar and elimination event times rFN_el, and the distances between
    environments D.
Set computational parameter elimination threshold ε.

▷ Create Pool of species                                                                  ▷ Algorithm 2
Pool ← PoolConstruction(B, C, Spread, B_min, B_max, C_min, C_max)

▷ Create interaction matrices for each environment.
▷ Govern effects of species on each other.
IntMatList ← ()
for i in 1 to N_env do
    IntMatList[i] ← InteractionConstruction(Pool, P)                                      ▷ Algorithm 3
end for
Reformat IntMatList using the matrix entries as blocks to form a single block diagonal matrix Mats.

▷ Create a function that describes the per-capita ecological/community dynamics            ▷ Algorithm 4
Dyn_e ← EcologicalConstruction(Pool[, 'Repr'], Mats, N_env)

▷ Identify a natural time scale                                                           ▷ Algorithm 5
λ ← CharacteristicRate(MatList).

▷ Build immigration/extirpation event history                                             ▷ Algorithm 6
E ← EventConstruction(B + C, N_env, N_ar, N_el, λ, λ, rFN_ar, rFN_el)

▷ Create a function that describes the movement of abundance between environments          ▷ Algorithm 7
Dyn_d ← SpatialConstruction(D, repeat((1), times = B + C))

▷ Combine features and run numerical integration for abundance densities through time
return PCA(Pool, N_env, E, Dyn_e, Dyn_d, ε)                                               ▷ Algorithm 1
```

---

Beyond the description of how the code works, we have also included the code used to generate example cases, used in Figure 3, and the code used to generate Figures 3 and 4 in the main text. These can be found in the subdirectories of experiments. To generate Figure 3, one sets their working directory to Figure3-ExampleOutcomes and first runs MNA-Image-ExampleOutcome-Create.R with parameters `PerIslandDistance <- 1` and `Space <- "Ring"`, `PerIslandDistance <-` $10^5$, and `PerIslandDistance <- Inf`. (Note that for the first run, one needs `CalculatePoolAndMatrices <- TRUE`, after which it can be set to false to avoid recalculating.) This generates three data sets (in a date based subdirectory) with file names dependent on their parameters: "MNA-ExampleExtProp-Result", number of environments, structure of patch connections, `PerIslandDistance`, immigration rate modifier, extirpation rate modifier, and the extirpation proportion. These data sets have the same pool, interaction matrices, and event histories, the first two of which are saved in "MNA-ExampleOutcome-PoolMats-Env10.RData". Then, one runs `MNA-Image-ExampleOutcome-Presence-Fig3.R`, although note that you may need to change the folder path on line 294 depending on the date on which you generate the data. Note that the "`by_for_thinning`" parameter will determine how much of the data is used for plotting. Lower values (minimum 1) should be more accurate, but much more time intensive. The plot is "MNA-Image-Example-Presence.png".

Finally, the data used for the main results shown in Figure 4 is generated with the following pipeline:

1. MNA-MasterAttempt-GenerateCases.R,

2. MNA-MasterAttempt-GeneratePMEsParallel.R (parallelised, one command line argument corresponding to the number of cores),
3. MNA-MasterAttempt-VikingRunHalves.R (parallelised, two command line arguments corresponding to case, A, B, or C, and parameter combination and replication, a number in 1 - 450, 1 - 180, and 1 - 1800 respectively),
4. Viking_HandleOutput_DiversityBC.R (parallelised, one command line argument corresponding to the number of cores),
5. Viking_HandleDiversity_Combine.R (parallelised, one command line argument corresponding to the number of cores),
6. Viking_HandleOutput_Invadability2Burnout.R (parallelised, one command line argument corresponding to the number of cores),
7. Viking_HandleInvadability_Combine.R (parallelised, one command line argument corresponding to the number of cores),
8. Viking_HandleOutput_TimeJaccard.R (parallelised, one command line argument corresponding to the number of cores),
9. Viking_HandleTimeJaccard_Combine.R (parallelised, one command line argument corresponding to the number of cores), and
10. MNA-Image-VikingOut-Figure4.R.

This, in order, creates the use cases, populates their starting parameters, runs each use case to generate the abundances through time and event results, processes the abundances to return diversity metrics, then aggregates the diversity metrics, repeats for invasibility and temporal Jaccard, and then plots Figure 4. Note that this is not a quick process, and the Viking Computing Cluster at the University of York was used to facilitate the process. All downstream random seeds are generated from the initial random seed when generating the cases to improve reproducibility. Note that, unlike in the example case, the pools, interaction matrices, and event histories are not necessarily exchangeable between runs. Each parameter combination has 10 pools, each with 10 associated interaction matrices and 10 histories, for a total goal of 100 runs for each parameter combination.

**C. Algorithms.** Algorithms in this section are as follows:

- Algorithm 1, Parallel Community Assembly (PCA), in which the primary function bundles together the constituent parts for the solver to use,
- Algorithm 2, PoolConstruction, in which the construction of the pool of species is detailed,
- Algorithm 3, InteractionConstruction, in which the construction of the interactions between abundance densities of species is detailed,
- Algorithm 4, EcologicalConstruction, in which the function that governs the dynamics of the ecosystem is constructed,
- Algorithm 5, CharacteristicRate, in which we describe how we approximate the characteristic rate of the ecological dynamics,
- Algorithm 6, EventConstruction, in which the construction of immigration and extirpation event histories is detailed, and
- Algorithm 7, SpatialConstruction, in which the function that governs the dynamics of inter-patch movement is described.

Algorithm 1 assumes that all of the other function outputs have been created. It takes these outputs and creates appropriate times to sample the abundance densities and the abundance time derivative function ($Dyn$). Note that the time steps must be “small” to make sure the results are smooth; jumping from event to event directly tends to result in numerical instability with steps that are too big and make the abundance densities appear to tend to infinity. We also set an initial abundance density of 0 for all species and run the LSODA algorithm.

Algorithm 2 then starts the pipeline that ends in Algorithm 1 by constructing our fixed list of species. This follows the work of Law and Morton 1996 by assigning to each species its trophic level, size (conditioned on trophic level), and reproduction rate (conditioned on trophic level and size).

Algorithm 3 is used to construct a matrix governing the interactions between species within a patch following Law and Morton’s 1996 work. We begin with an “empty” square real matrix and will populate each of its entries according to intraspecific interactions (the diagonal) or its predator-prey interactions (off-diagonal). Predator-prey interactions are size-structured here, with larger things eating smaller things, but having a preference determined by the parameters. Energy flow is reduced by a factor to reflect that not all energy is efficiently converted by the predator. Each entry of the matrix is initially calculated for the average interaction between two species, with the last step being to add Gaussian noise. This noise is what makes each patch different from the others; species will have the same type of interactions and the interactions will obey roughly the same ecology, but the exact values will differ to reflect (minor) differences in environment.

Algorithm 4 specifies our generalised Lotka-Volterra dynamics as reproduction added to interactions on the per-capita scale.

Algorithm 5 reflects how we try to obtain a measure of the time scale upon which our interactions occur. An immediate problem is that we do not have knowledge of what non-trivial equilibrium the assembly system might settle into, especially as we do not require feasibility of any non-trivial set of species. Our solution is to consider the time scale of the (whole) interaction matrix itself, rather than of the dynamical system. As the structure of the interaction matrix of the system will be block diagonal (species only interact with each other locally in this model) and thus we need only compute the eigenvalues of the blocks individually. Since we are only interested in the time scale, rather than the stability (which will emerge naturally for any of our obtained systems due to the structure of the system), we specifically take the largest magnitude of the eigenvalues as our characteristic rate.

Algorithm 6 details how we construct the neutral assembly sequence. Immigration and extirpation events are created independently of each other but in an analogous fashion: sample waiting times, sample species targets, sample environment

**Algorithm 1** Parallel Community Assembly (PCA)

[This is the "main" function that processes the outputs of the other functions to provide to LSODA, which performs the actual numerical integration. This corresponds to "MultipleNumericalAssembly_Dispersal" in the R code. ]

**function** PCA($Pool, N_{env}, E, Dyn_e, Dyn_d, \epsilon$)

  $Pool$: DF ▷ pool of species

  $N_{env}$: Nat ▷ number of environments

  $E$: DF($Time, Type, Species, Environment$) ▷ events

  $Dyn_e$: Func($Population, \ldots$) ▷ takes population; returns 1D-Object of per capita net gains by species and location, see Algorithm 4

  $Dyn_d$: Func($Population, \ldots$) ▷ takes population; returns 1D-Object of net dispersal by species and location, see Algorithm 7

  $\epsilon$: $\text{Real}_{>0}$ ▷ threshold for zeroing of a population

**Require:** $E$ sorted by $Time$

  $Dyn \leftarrow \text{Func}(Popul)\{Popul \times Dyn_e(Popul) + Dyn_d(Popul)\}$

  ▷ Construct set of times to be returned from solver;

  ▷ will need 0 and Event Times for interpolation.

  $T \leftarrow \text{cat}(0, E[, `Time'])$

  $T_{out} \leftarrow$ 1D-Object of finer grain interpolated times $T$

  ▷ Run ODE solver to get 2D-Object of abundance densities over time.

  $y \leftarrow \text{repeat}((0), \text{times} = \text{rows}(Pool) \times N_{env})$ ▷ Initial pop. of zeros

  $AbundOverTime \leftarrow \text{LSODA}(y, T_{out}, Dyn, E, \epsilon)$

  **return** $AbundOverTime$

**end function**

 

**Algorithm 2** Pool Construction

[Generate the species pool by taking in the requested number of species for the trophic levels and randomly generating their two traits of size and reproductive rate. This corresponds to "LawMorton1996_species" in the R code.]

**function** PoolConstruction($B, C, P, B_{\min}, B_{\max}, C_{\min}, C_{\max}$)
    $B$: Nat ▷ Number of basal species
    $C$: Nat ▷ Number of consumer species
    $Spread$: $\text{Real}_{>0}$ ▷ Ratio of standard deviation to mean, default 0.1
    $B_{\min}$: $\text{Real}_{>0}$ ▷ Minimum basal body size, default $10^{-2}$
    $B_{\max}$: $\text{Real}_{>0}$ ▷ Maximum basal body size, default $10^{-1}$
    $C_{\min}$: $\text{Real}_{>0}$ ▷ Minimum consumer body size, default $10^{-1}$
    $C_{\max}$: $\text{Real}_{>0}$ ▷ Maximum consumer body size, default $10^{0}$
    $ID \leftarrow (1, \ldots, B + C)$
    $Type \leftarrow \text{cat}(\text{repeat}((\text{"Basal"}), \text{times} = B), \text{repeat}((\text{"Consumer"}), \text{times} = C))$
    ▷ Select size unif. at rand. over $\log_{10}$ scales
    $Size \leftarrow \text{cat}(\text{runif}(\text{times} = B, \min = \log_{10} B_{\min}, \max = \log_{10} B_{\max}), \text{runif}(\text{times} = C, \min = \log_{10} C_{\min}, \max = \log_{10} C_{\max}))$
    ▷ Get mean reproduction rates
    $Repr \leftarrow \text{cat}(10^{-1-0.25\times Size[1:B]}, \text{repeat}((-0.1), \text{times} = C))$
    ▷ Add noise from a truncated at 0 normal distribution
    $Repr \leftarrow \text{sign}(Repr) \times \text{rnorm0trunc}(|Repr|, |Repr| Spread)$
    $Size \leftarrow 10^{Size}$ ▷ Convert to real sizes
    $Pool \leftarrow \text{DF}(ID, Type, Size, Repr)$
    **return** $Pool$
**end function**

**Algorithm 3** Interaction Matrix Construction

[Generate an interaction matrix by comparing species and assigning interactions by trophic level and size. This follows Law and Morton's 1996 appendix closely. This corresponds to "LawMorton1996_CommunityMat" and "LawMorton1996_aij" in the R code.]

**function** InteractionConstruction($Pool, Strg, Pref, Spec, Effi, EqDn$)
    $Pool$: DF($ID, Type, Size, Repr, \ldots$) ▷ species data frame from Alg.2
    $Strg$: $\text{Real}_{>0}$ ▷ Interaction strength, default 0.01
    $Pref$: $\text{Real}_{>0}$ ▷ Consumer body size preference, default 10
    $Spec$: $\text{Real}_{>0}$ ▷ Consumer prey nonspecificity, default 0.5
    $Effi$: $\text{Real}_{>0}$ ▷ Predation energy efficiency, default 0.2
    $EqDn$: $\text{Real}_{>0}$ ▷ Basal equilibrium biomass density, default 100
    $Spread$: $\text{Real}_{>0}$ ▷ Ratio of standard deviation to mean, default 0.1
  $Rate \leftarrow$ Zero Square Real $\text{Matrix}_{\text{rows}(Pool)}$ ▷ Initialised to all 0

  ▷ Diagonal: Basal intraspecific impact (no intraspecific Consumer)
  **for** $i$ in 1 to rows($Pool$) **do**
    **if** $Pool[i, \text{'}Type\text{'}] =$ "Basal" **then**
      ▷ Set basal to same equilibrium biomass
      $Rate[i, i] \leftarrow -Pool[i, \text{'}Repr\text{'}] \times Pool[i, \text{'}Size\text{'}]/EqDn$
    **end if**
  **end for**

  ▷ Interspecific impact, species $j$ on species $i$ (no Basal/Basal)
  **for** $i$ in 1 to rows($Pool$) **do**
    **for** $j$ in 1 to rows($Pool$) **do**
      **if** $Pool[i, \text{'}Size\text{'}] < Pool[j, \text{'}Size\text{'}]$
          & $Pool[j, \text{'}Type\text{'}] =$ "Consumer" **then**
        ▷ Decrease in prey $i$ due to larger consumer $j$
        $Rate[i, j] \leftarrow -Strg \exp\left(-\left(\log_{10}\left(Pref\frac{Pool[i, \text{'}Size\text{'}]}{Pool[j, \text{'}Size\text{'}]}\right)/Spec\right)^2\right)$
      **else if** $Pool[i, \text{'}Size\text{'}] > Pool[j, \text{'}Size\text{'}]$
          & $Pool[i, \text{'}Type\text{'}] =$ "Consumer" **then**
        ▷ Increase in larger consumer $i$ from prey $j$
        ▷ Calculate $Rate[j, i]$, then apply efficiency/conversion
        $R \leftarrow -Strg \exp\left(-\left(\log_{10}\left(Pref\frac{Pool[j, \text{'}Size\text{'}]}{Pool[i, \text{'}Size\text{'}]}\right)/Spec\right)^2\right)$
        $Rate[i, j] \leftarrow -R \times Effi\frac{Pool[j, \text{'}Size\text{'}]}{Pool[i, \text{'}Size\text{'}]}$
      **end if**
    **end for**
  **end for**

  ▷ Add noise (to non-zero rates)
  **for** $i$ in 1 to rows($Pool$) **do**
    **for** $j$ in 1 to rows($Pool$) **do**
      $R \leftarrow Rate[i, j]$
      $Rate[i, j] \leftarrow \text{sign}(R) \times \text{rnorm0trunc}(|R|, |R|Spread)$
    **end for**
  **end for**

  **return** $Rate$
**end function**

**Algorithm 4** Construct per capita ecological dynamics

[Follows generalised Lotka-Volterra dynamics using the interaction matrix. The output is the lambda function which will handle abundance densities (populations) across environments and return their (per-capita) net growth rates. This corresponds to “PerCapitaDynamics_Type1” in the R code.]

**function** ECOLOGICALCONSTRUCTION($Repr, Mat, N_{env}$)
  $Repr$: 1D-Object of Reals ▷ reproduction rates of species
  $Mat$: 2D-Object of Reals ▷ overall species interaction matrix
  $N_{env}$: Nat ▷ number of environments
  $R \leftarrow \text{repeat}(Repr, \text{times} = N_{env})$
  $Dyn_e \leftarrow \text{Func}(y, \ldots)\{R + Mat \times y\}$ ▷ Matrix vector multiplication
  **return** $Dyn_e$
**end function**

**Algorithm 5** Identify approximate characteristic rate.

[In principle, this should be the eigenvalues of the system near equilibrium. Since we do not know the equilibrium ahead of time, we instead use the characteristic rate of the interactions.]

**function** CHARACTERISTICRATE($MatList$)
  $MatList$: List of Real Matrices ▷ Each environment’s interaction matrix
  $\lambda \leftarrow \langle\rangle$
  **for** $Mat$ in $MatList$ **do**
    $\lambda \leftarrow \text{cat}(\lambda, |\text{eigenvalues}(Mat)|)$
  **end for**
  **return** $\max(\lambda)$
**end function**

**Algorithm 6** Construct the assembly sequence events

[Creates a “history” of events by randomly sampling times, species and environments before combining them. This corresponds to “CreateAssemblySequence” in the R code.]

**function** EVENTCONSTRUCTION($N_{sp}, N_{env}, N_{ar}, N_{el}, Rate_{ar}, Rate_{el}, rFN_{ar}, rFN_{el}$)
  $N_{sp}$: Nat ▷ Number of species
  $N_{env}$: Nat ▷ Number of environments
  $N_{ar}$: Nat ▷ Number of arrival/immigration events
  $N_{el}$: Nat ▷ Number of elimination/extirpation events
  $Rate_{ar}$: $\text{Real}_{>0}$ ▷ Rate of arrival/immigration events
  $Rate_{el}$: $\text{Real}_{>0}$ ▷ Rate of elimination/extirpation events
  $rFN_{ar}$: Function: Nat $\times$ $\text{Real}_{>0} \rightarrow \text{Real}_{>0}$ Array ▷ Function to sample arrival/immigration events
  $rFN_{el}$: Function: Nat $\times$ $\text{Real}_{>0} \rightarrow \text{Real}_{>0}$ Array ▷ Function to sample elimination/extirpation events
  $Ar \leftarrow rFN_{ar}(N_{ar}, Rate_{ar})$ ▷ Arrival times
  $El \leftarrow rFN_{el}(N_{el}, Rate_{el})$ ▷ Elimination times
  $Species \leftarrow \text{rpick}(\text{times} = \text{length}(Ar) + \text{length}(El), \text{min} = 1, \text{max} = N_{sp})$
  $Envir \leftarrow \text{rpick}(\text{times} = \text{length}(Ar) + \text{length}(El), \text{min} = 1, \text{max} = N_{env})$
  $E \leftarrow \text{DF}(\text{cat}(Ar, El), Species, Envir)$ ▷ Combine
  $E \leftarrow E[\text{sortedOrder}(\text{cat}(Ar, El)), ]$ ▷ Sort data frame by event times
  **return** $E$
**end function**

targets, and then combine into a single “history”.

Algorithm 7 details how we take an adjacency matrix describing the connections between patches and the ability of species to travel and create a single matrix of abundance transfer rates. The premise is straightforward: the adjacency matrix describes distance from location to location and the species have “speeds” (set to 1 in our work here). The rate of travel for a species moving from one location to the next is then found by multiplication and these form the diagonal of a matrix block. (Other entries are 0; species cannot change from one type to another by moving patches.) These blocks can then be assembled into the dispersal matrix. Then, to impose mass conservation, we set the diagonal of the dispersal matrix (the “amount of travel from a patch to itself”) to be the loss due to all travel away from that patch. This is then formulated into a function of the abundance, for use within the LSODA dynamics.

**Algorithm 7** Construct spatial dynamics

[Create a banded matrix from diagonal block matrices that describe how the species (within block) move from one patch to the next (position of block in full matrix). This corresponds to “CreateDispersalMatrix” in the R code.]

**function** SPATIALCONSTRUCTION($D, S$)
  ▷ Adjacency matrix of environment-environment distances graph
  $D$: 2D-Object of $\text{Real}_{>0}$
  ▷ Speeds of individual species in the pool; default repeat((1),rows($Pool$))
  $S$: 1D-Object of $\text{Real}_{>0}$

  ▷ Combine spatial layout with species speeds
  $Mat \leftarrow$ Square $\text{Real}_{\geq 0}$ $\text{Matrix}_{\text{length}(S)\times\text{rows}(D)}$
  ▷ Mat acts as a matrix with rows($D$) square blocks of length($S$) rows.
  ▷ $\text{Block}_{r,c}$'s diagonal describes the abundance that moves to $r$ from $c$.
  ▷ As a matrix, elements are initialised to 0.
  ▷ Travel from $c$ to $r$ is $Mat[r, c]$
  ▷ Total travel from $r$ is $Mat[r, r]$
  ▷ Travel is in blocks and bands since species cannot change identity
  $Size \leftarrow \text{length}(S)$
  ▷ Iterate over blocks:
  **for** $r$ in 1 to rows($D$) **do**
    **for** $c$ in 1 to cols($D$) **do**
      **if** $r = c$ **then next** ▷ ignore diagonal blocks for now
      **end if**
      $Diag \leftarrow S/D[r, c]$ ▷ Array of rates to block's diagonal
      $Blkx \leftarrow (r-1) \times Size + (1, \ldots, Size)$ ▷ Row coordinates of block
      $Blky \leftarrow (c-1) \times Size + (1, \ldots, Size)$ ▷ Col. coordinates of block
      diagonal($Mat[Blkx, Blky]$) $\leftarrow Diag$ ▷ Add block to main matrix
    **end for**
  **end for**
  ▷ Mass conservation by deducting travel along the full diagonal.
  diagonal($Mat$) $\leftarrow -$columnSums($Mat$)
  $Dyn_d \leftarrow \text{Func}(Popul, \ldots)\{Mat \times Popul\}$
  **return** $Dyn_d$
**end function**

#### 6. References, Tables, and Images


**Table S1. System parameters that are varied between simulations.**

| Fixed Parameter | Value | Notes |
|---|---|---|
| Pools per Parameter Set | 10 | |
| Histories per Pool | 10 | |
| Islands per Pool | 10 | |
| Elimination Threshold ($\epsilon$) | $10^{-4}$ | Numerical Threshold |
| Establishment Density | $4 \times 10^{3}\epsilon = 0.4$ | See (4) |
| Basal Species | 34 | |
| Consumer Species | 66 | |
| Species ‘speeds’ | 1 | |
| Interaction Strength ($k_1$) | 0.01 | See (2) |
| Preferred Prey Size ($k_2$) | 10 | See (2) |
| Willingness to Deviate ($k_3$) | 0.5 | See (2) |
| Conversion Efficiency ($k_4$) | 0.2 | See (2) |
| Basal Equilibrium Biomass ($k_5$) | 100 | See (2) |
| Noise Strength ($k_6$) | 0.1 | See (2) |

**Table S2. System parameters that are varied between simulations.**

| Experimental Parameter | Value | Notes |
|---|---|---|
| Basal Min. Size | $-3, -2$ | $\log_{10}$ scale |
| Basal Max. Size | $-1$ | $\log_{10}$ scale |
| Consumer Min. Size | $-1$ | $\log_{10}$ scale |
| Consumer Max. Size | $0, 1$ | $\log_{10}$ scale |
| Inter-patch Variation | 0, 1, 2 | Multiplies $k_6$.[1] |
| Neutral Immigration | $0.1, 1, 10$ | Multiplier.[2] |
| Neutral Extirpation | $0, 0.1, 1, 10$ | Multiplier.[2] |
| Inter-patch Resistance | $10^{0}$ to $10^{9}$, Infinite | Distance units. |
| Inter-patch Dispersal | 86.47% to $2 \times 10^{-8}$%, 0% | [3] |

See Table S3 for more details on combinations of parameters.

(1) Multiplies $k_6$ only when calculating the interaction matrix. This multiplier does not affect the species pool.

(2) Neutral parameters are multipliers on the characteristic rate.

(3) Inter-patch dispersal is equivalent to distance and is measured as the (theoretical) percentage abundance density turnover per unit time due to dispersal.

**Table S3. Sets of Simulations Conducted.**

| Set | Pool[1] | Var.[6] | (Imm., Ext.)[7] | Simulations | Completed |
|---|---|---|---|---|---|
| A | Base Case[2] | 1 | (1, 0)<br>(0.1, 0.1)<br>(0.1, 10)<br>(10, 0.1)<br>(10, 10) | 5500 | 2828 |
| B | Base Case[2] | 0 | (1, 1)<br>(1, 0) | 2200 | 2200 |
| C | Base Case[2]<br>Smaller Basal[3]<br>Larger Consumer[4]<br>Smaller and Larger[5] | 1 | (1, 1)<br>(1, 10)<br>(1, 0.1)<br>(10, 1)<br>(0.1, 1) | 22000 | 18042 |
| D | Base Case[2] | 2 | (1, 1) | 1100 | 1100 |

All above sets were tested with each inter-patch resistance: from $10^0$ to $10^9$ at 10 even intervals on logarithmic scale and an infinite amount of resistance. Additionally, there was a fifth set of 3742 (of 4600) simulations, composed of combinations not described above of set A's (Imm., Ext.) and pools (-3, -1, -1, 0), (-2, -1, -1, 1), and (-3, -1, -1, 1) with coefficient of variation $k_6 = 0.1$ and pool (-2, -1, -1, 0) with $k_6 = 0$. This fifth set was only tested on dispersal rates of 0.0198 and 0.1813 to ensure robustness of the trends at these values. This is apparent in Figure S1 and Figure S2.

(1) Pool refers to the (log-scale) body size ranges, which can be characterised by basal minimum, basal maximum, consumer minimum, and consumer maximum, as shown with Table S2.

(2) Characteristic values (-2, -1, -1, 0).

(3) Characteristic values (-3, -1, -1, 0).

(4) Characteristic values (-2, -1, -1, 1).

(5) Characteristic values (-3, -1, -1, 1).

(6) Var. refers to the multiplier on $k_6$ for inter-patch variation of the interaction matrix (but not the initial pool). Set B refers to the homogeneous patch environments case. Set D refers to the highly heterogeneous patch environments case.

(7) Imm. and Ext. refer to immigration and extirpation event rate multipliers respectively.

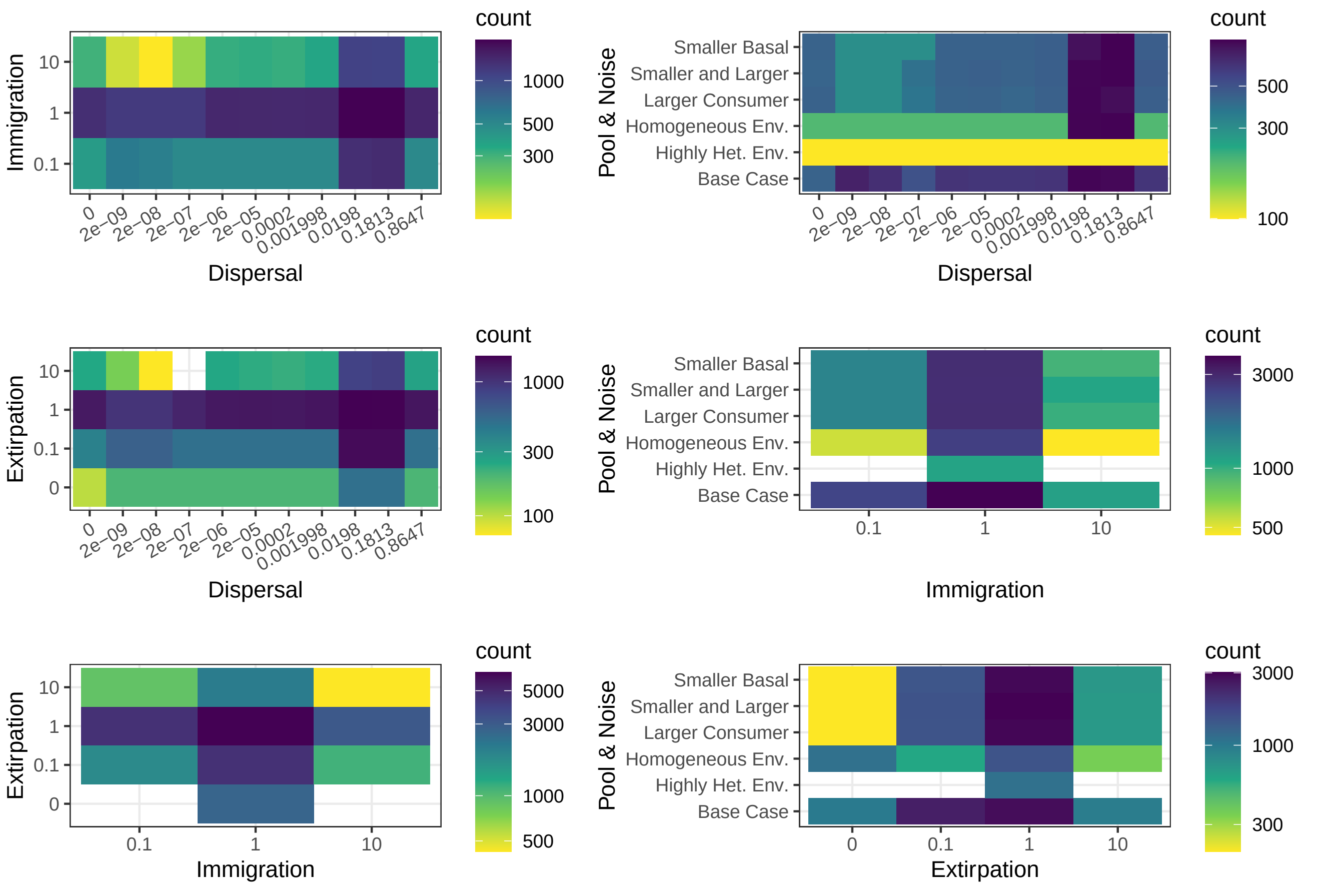


**Fig. S1.** Heat maps of the raw number of simulations completed with each parameter combination. All plots represent the same underlying set of simulations and numbers are just visualised for different combinations of parameters. Note that some combinations have very few simulations desired and no simulations were attempted for extirpation rate 0 and immigration rate 0.1 or for extirpation rate 0 and immigration rate 10. See Figure S2 for percentages of simulations completed.
Immigration and Extirpation are multipliers applied to the characteristic rate of the matrix in order to determine the overall rate of local, neutral species arrivals and removals. Dispersal is the proportion of abundance that can move to other patches in a single time unit. Pool & Noise refers to how the pool range limits and interaction matrix noise are varied. The cases, from bottom to top, refer to the base case (heterogeneous environments in the main text), to highly heterogeneous environments (i.e., doubled coefficient of variation when generating the interaction matrix), to homogeneous environments (i.e., no noise in the interaction matrix), to larger consumer species (i.e., an increase of 1 to the maximum logarithmic body size), to larger consumer species and smaller basal species (i.e., a decrease of 1 to the minimum logarithmic body size), and to smaller basal species.

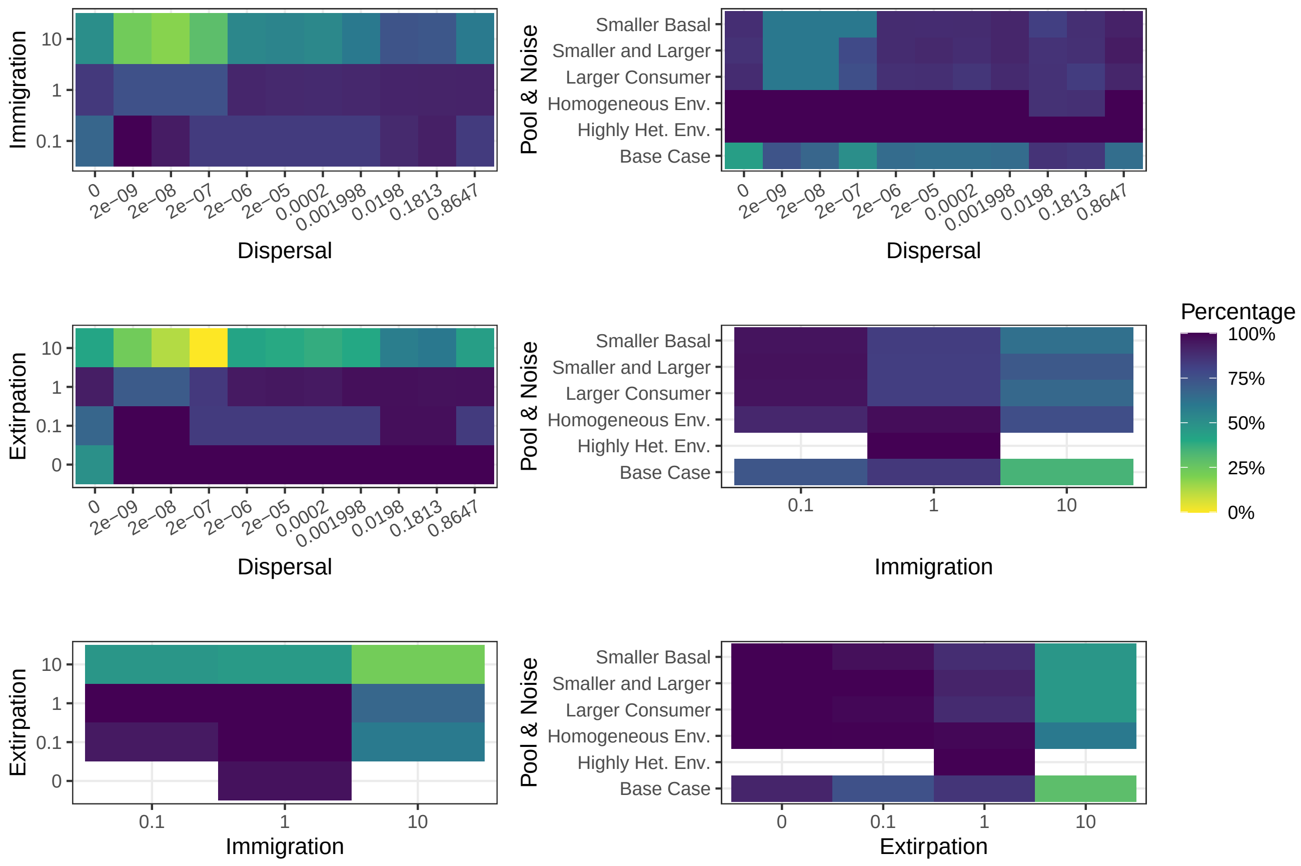


**Fig. S2.** Heat maps of the percentage of simulations completed for each parameter combination. Plots and values are otherwise as in Figure S1. No simulations were attempted for extirpation rate 0 and immigration rate 0.1 or for extirpation rate 0 and immigration rate 10.

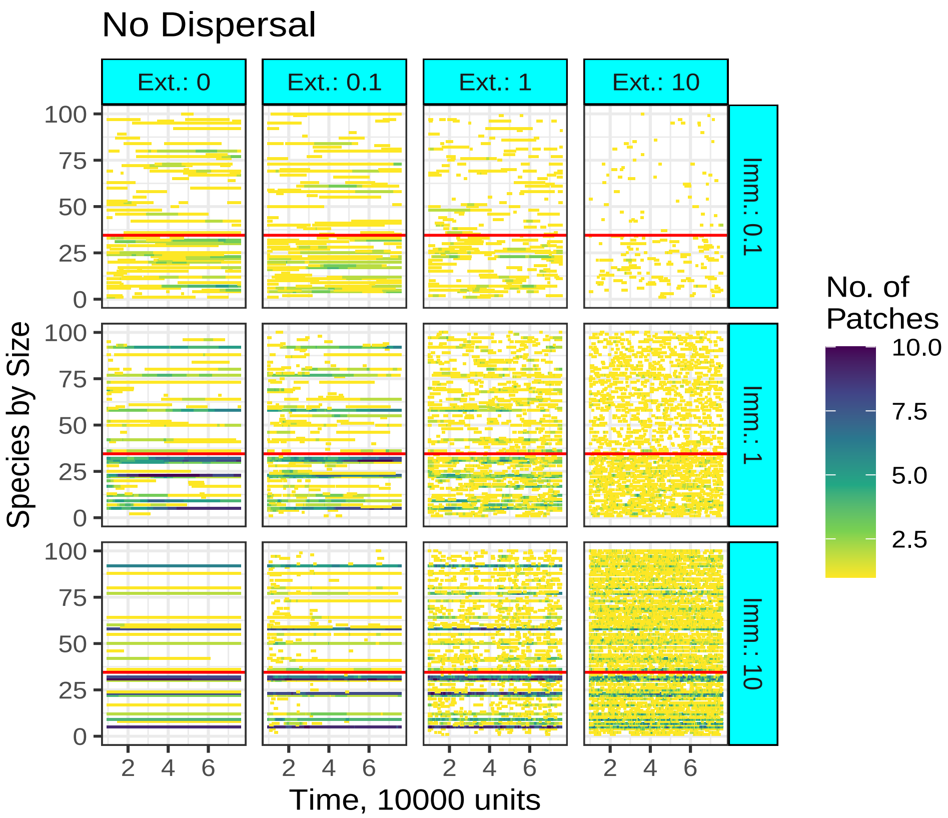


**Fig. S3.** Results for twelve simulations conducted with no dispersal between patches. As in Figure 3 (a) of the main text, we show the number of patches that each species, ordered by size, is present on, with basal species below the red line and consumer species above it for the no dispersal case. Note that each plot within the figure has its own history of immigration and extirpation events, although these histories are shared with Figure S4 and Figure S5. Event histories are random samples from all possible event histories (with these immigration and extirpation rates).

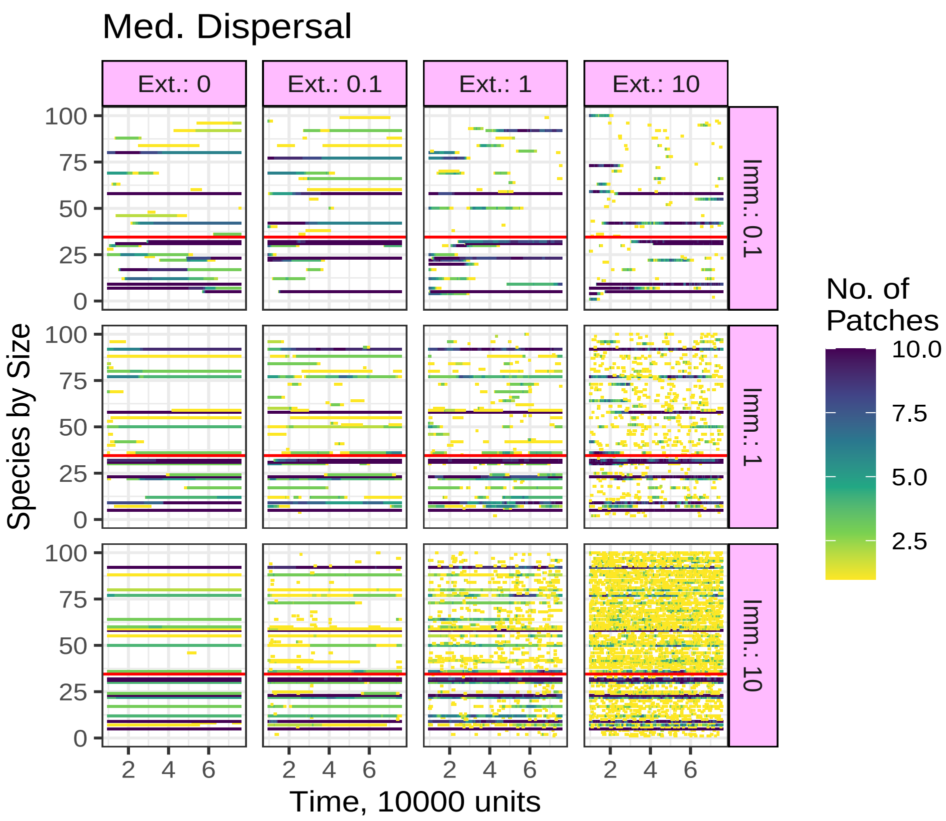


**Fig. S4.** As in Figure S3, but simulations instead have a medium amount of dispersal, equivalent to 2e-05 ($2 \times 10^{-5}$) abundance proportion moving in 1 time unit elsewhere.

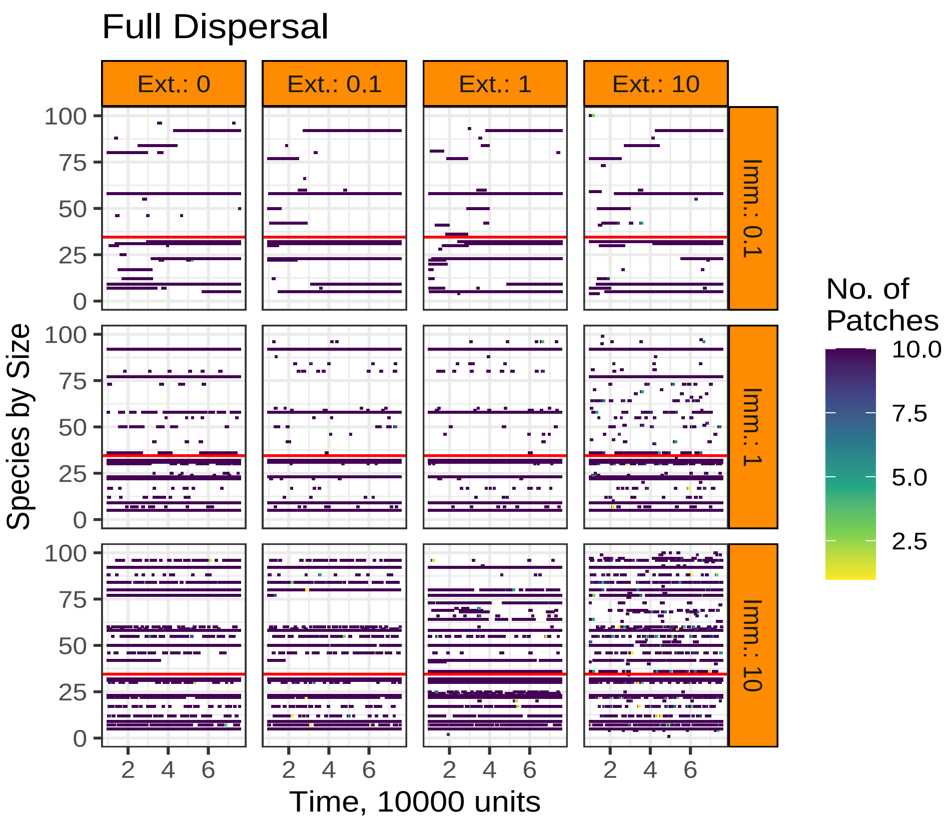


**Fig. S5.** As in Figure S3, but simulations instead have full dispersal, equivalent to 0.8647 abundance proportion moving in 1 time unit elsewhere.

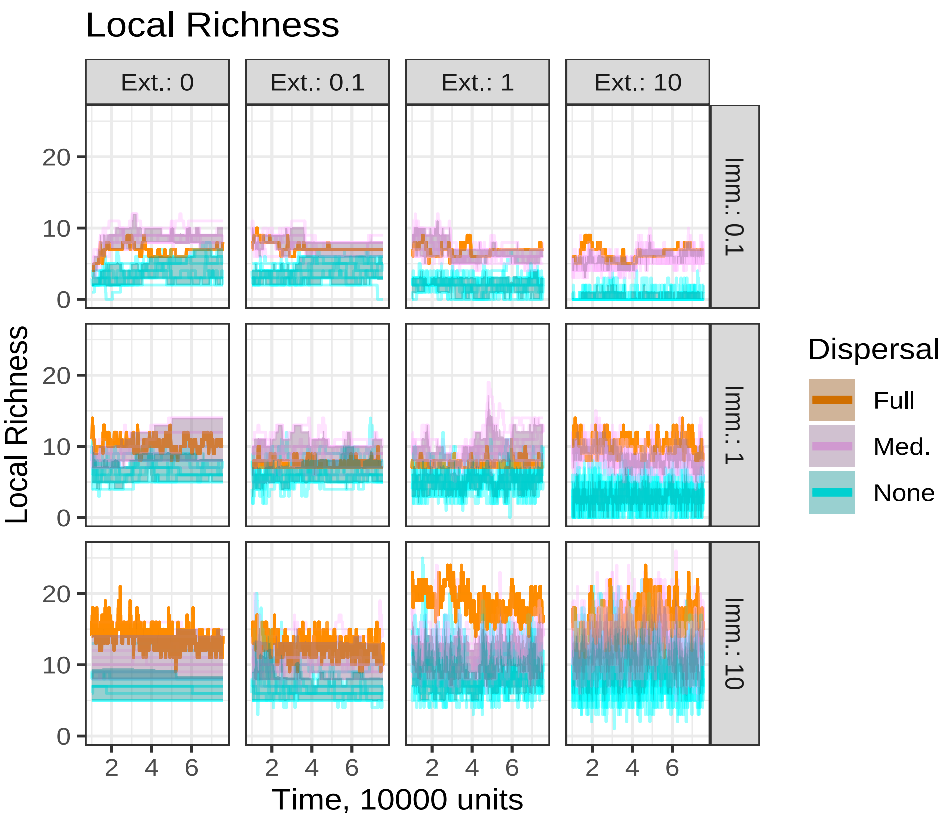


**Fig. S6.** Results for twelve simulations conducted with no dispersal between patches. As in Figure 3 (b) of the main text, we compare the local richness (number of species per patch through time) between the three cases in Figure S3, Figure S4, and Figure S5. Regional richness is examined in Figure S7 and spatial Jaccard distance is examined in Figure S8. The dark regions correspond to $80\%$ intervals. Individual lines correspond to individual patches. The results are created from event histories that are random samples from all possible event histories (with these immigration and extirpation rates).

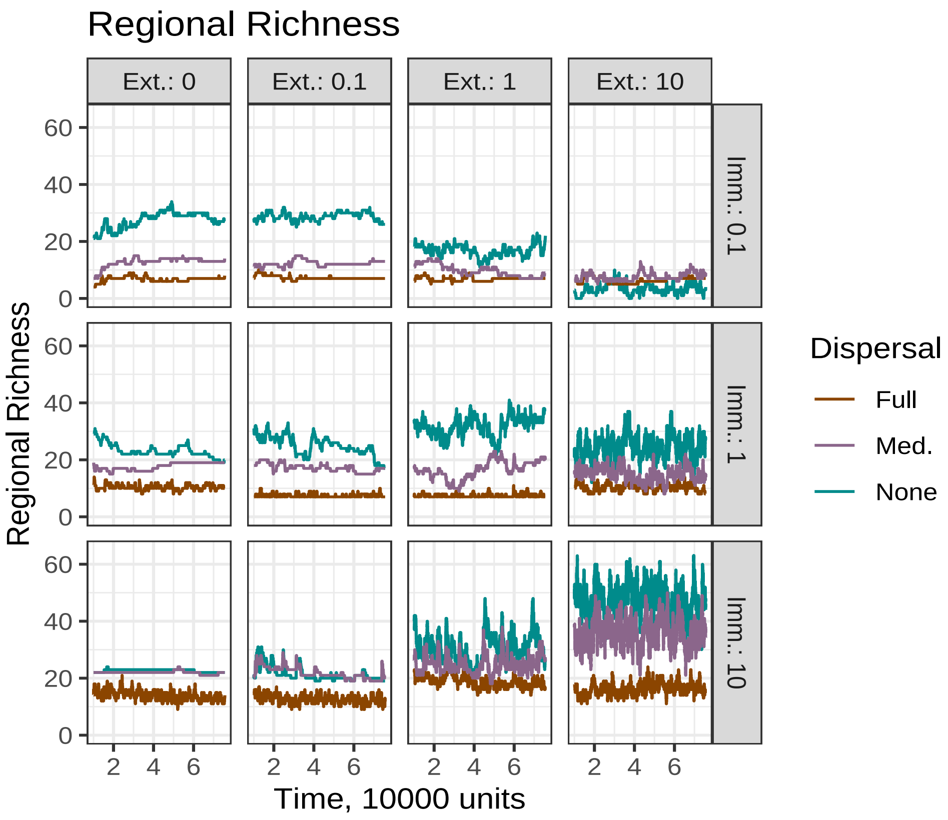


**Fig. S7.** As in Figure S6, but comparing regional richness of the simulations over time instead. In contrast to Figure S6 and Figure S8, there is only a single value for the entire metacommunity.

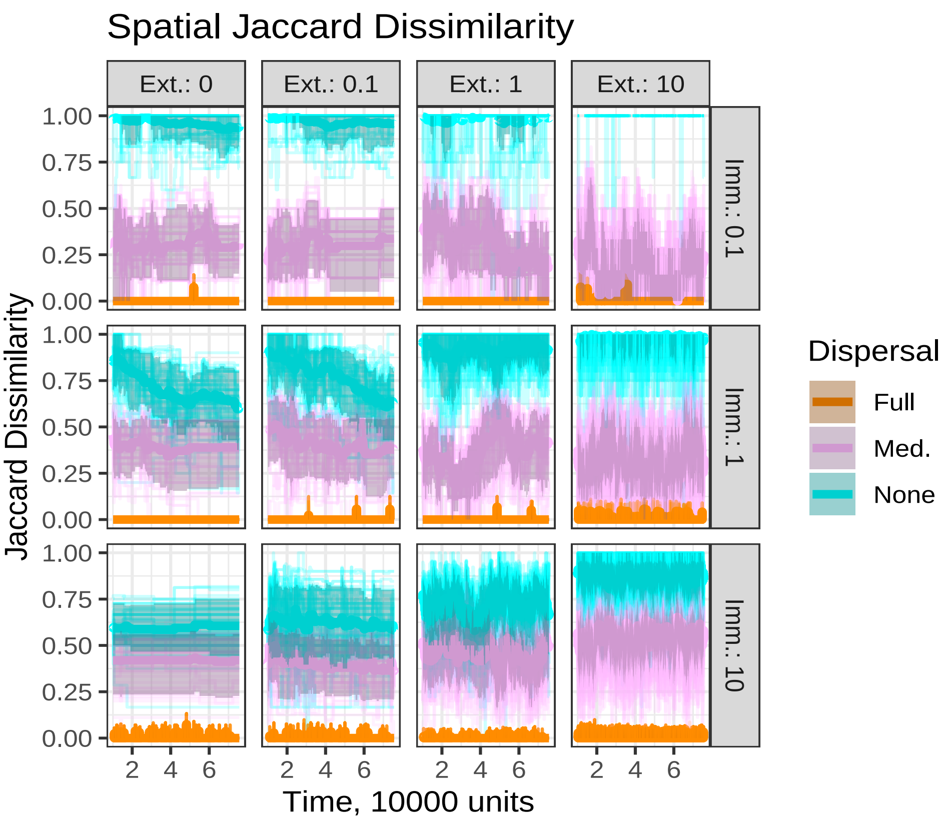


**Fig. S8.** As in Figure S6, but comparing pairwise (spatial) Jaccard distances of the simulations over time instead. The dark regions correspond to $80\%$ intervals.

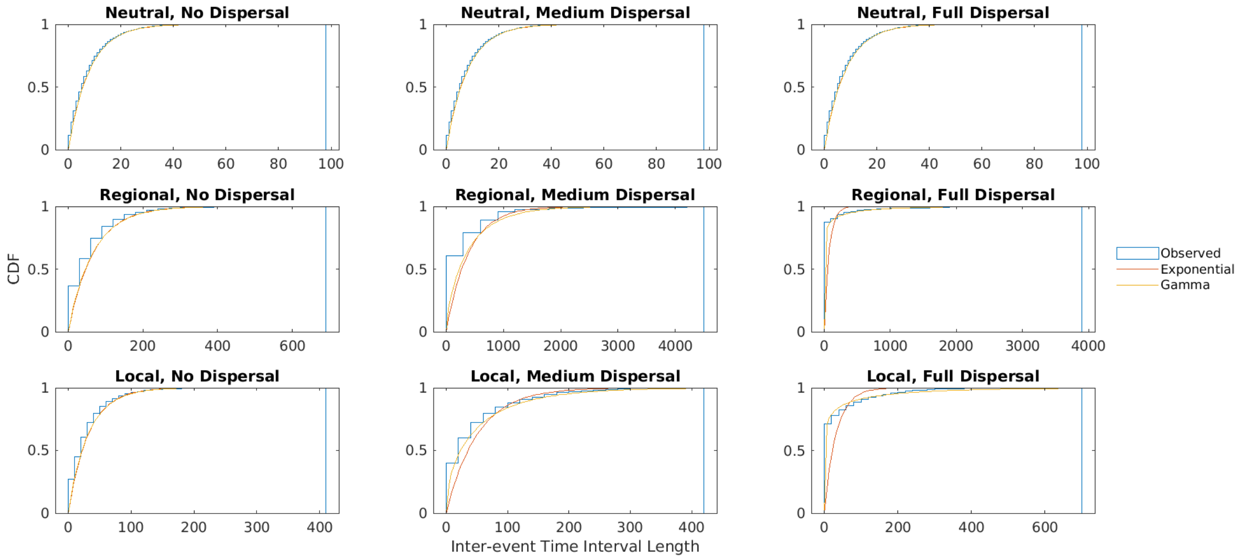


**Fig. S9.** We perform maximum likelihood estimation and fitting for exponential and gamma distributions to the immigration and extirpation events shown in Figure 3 (a), broken down by whether they are neutral immigration or extirpation events ("Neutral" events, including immigration and extirpation events that fail to establish a species on or remove a species from a patch), events at the archipelago scale that are either neutral or driven by community dynamics ("Regional" events, marking the immigration or extirpation of a species to or from all islands), or events at the island scale that are either neutral or driven by community dynamics (all "Local" events, marking the immigration or extirpation of a species to or from any island). Note the differing x-axis scales reflecting different observed maximum waiting times. The y-axis corresponds to the probability that the inter-event time interval ("waiting time") will be shorter than a given x-value.

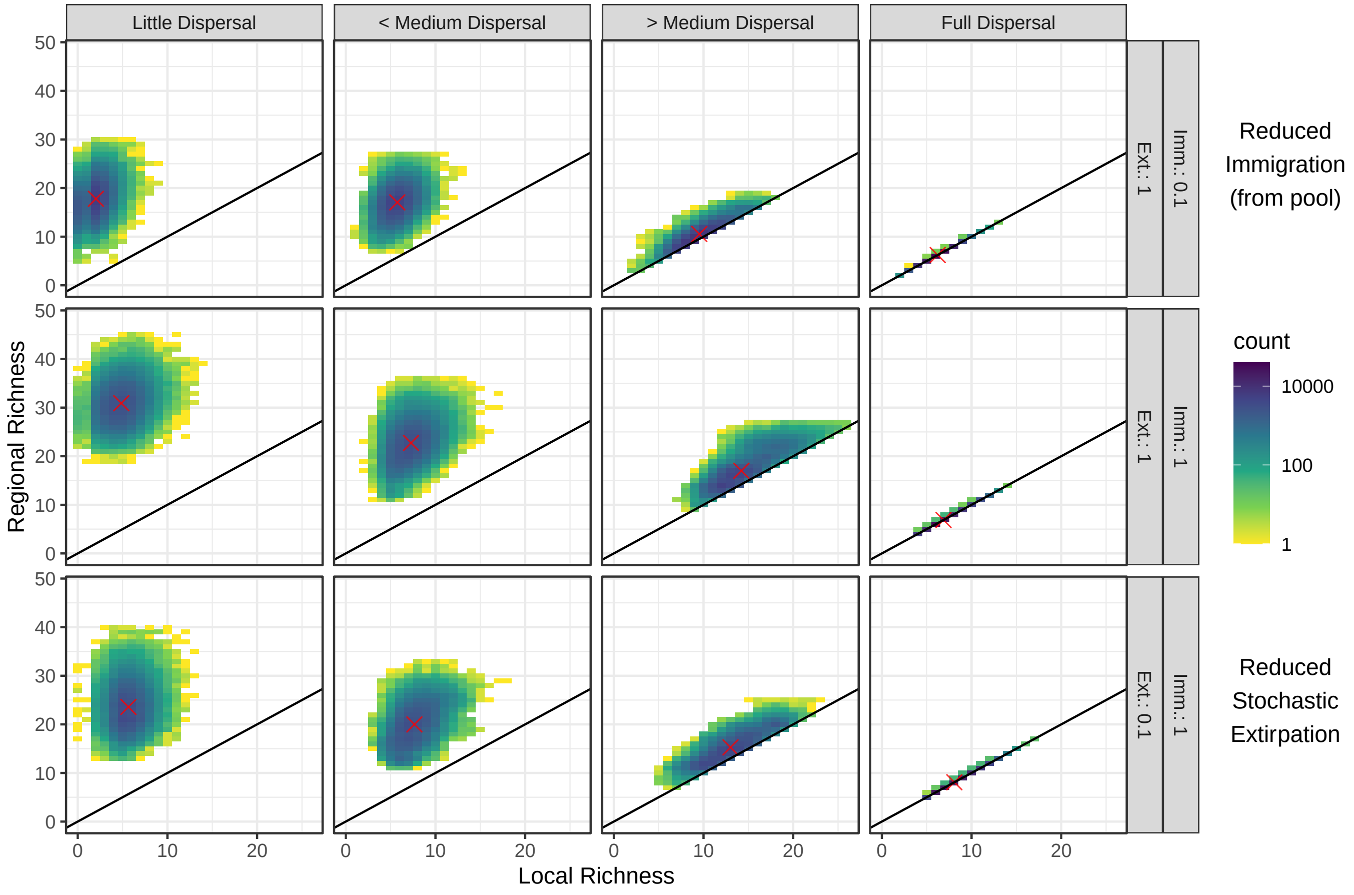


**Fig. S10.** We show heat maps of the space of local richness (alpha diversity) and regional richness (gamma diversity) values explored by the simulations between burn-in $(10^4)$ and burn-out $(6.5 \times 10^4)$ times. The number of times the patches in simulations visit the combination of richness values is counted, with darker regions (on a logarithmic scale) visited more often. Mean values are shown with red crosses. The black line is the physical local richness equal to regional richness line; no patch can have more species than there are species in the system. Not shown is the richness equal to pool richness boundary lines, reflecting the pool size (here, 100 on both axes). Dispersal increases from left to right, proceeding as little dispersal $(2 \times 10^{-9})$, $<$ medium dispersal $(2 \times 10^{-6})$, $>$ medium dispersal $(2 \times 10^{-3})$, and full dispersal $(0.8647)$ in order to show more of the evolution of the region than using just no, medium, and full would allow. The top row corresponds to a reduced immigration scenario, the middle row to the base case (equal stochastic immigration and stochastic extirpation), and the bottom row to a reduced stochastic extirpation scenario.

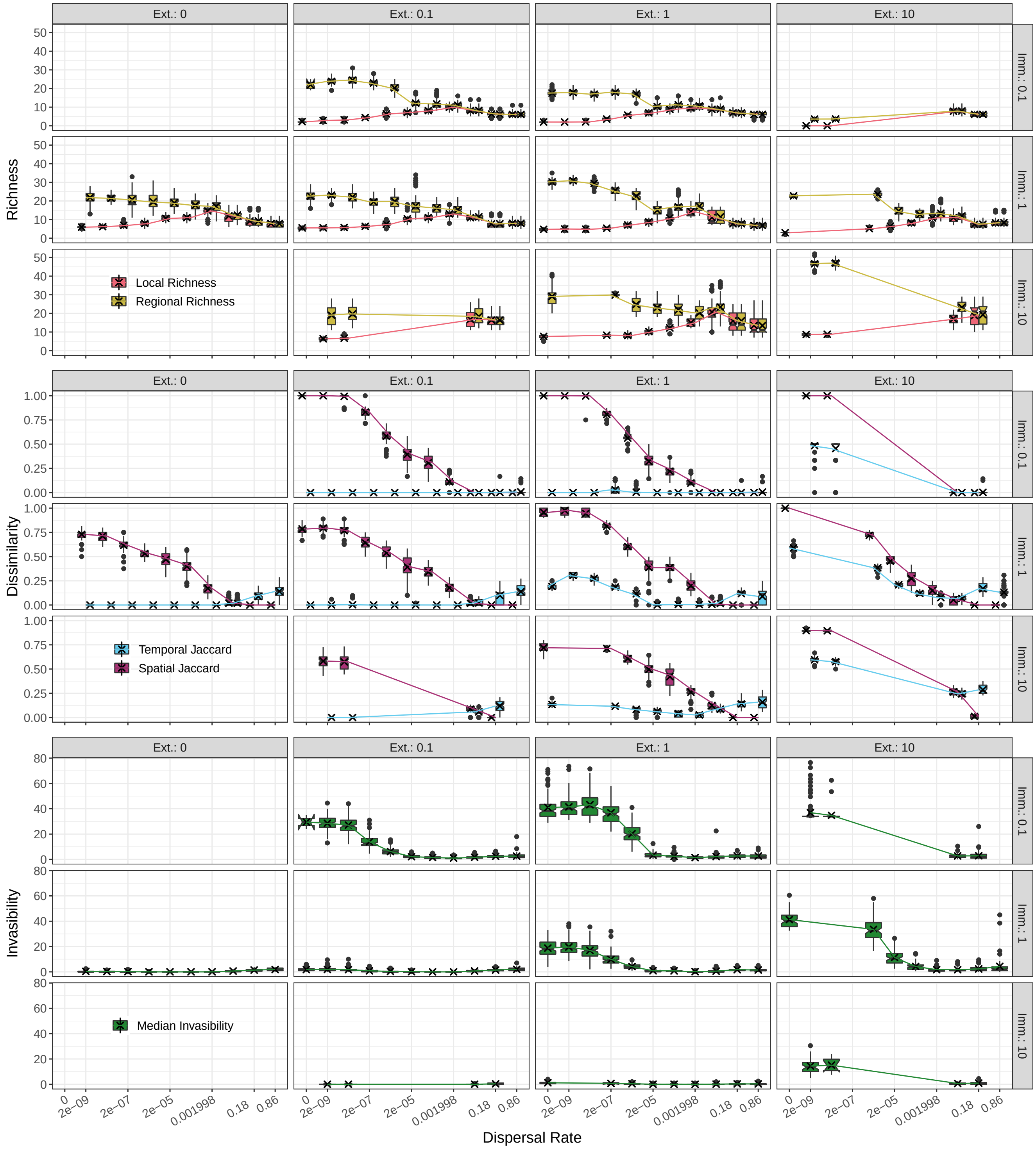


**Fig. S11.** As in Figure 4 of the main text, we consider five summary statistics of our simulations: median local richness (top, red), median regional richness (top, yellow), median temporal Jaccard dissimilarity (blue, middle), median spatial Jaccard dissimilarity (magenta, middle) and median end of simulation local invasibility (bottom, green). Simulations correspond to our base case with coefficient of variation of interaction matrices set to $k_6 = 0.1$, and we vary the immigration (Imm., rows) and extirpation (Ext., columns) rate multipliers. Trends generally follow those described in the main text (see also Imm.: 1, 2nd row, and Ext.: 1, 3rd column), except as noted in Section 4. Note that an extirpation multiplier of 0 removes extirpation events from the simulation. In cases where dispersal rates do not have corresponding box plots, no "base case" simulations were conducted, see Table S3.

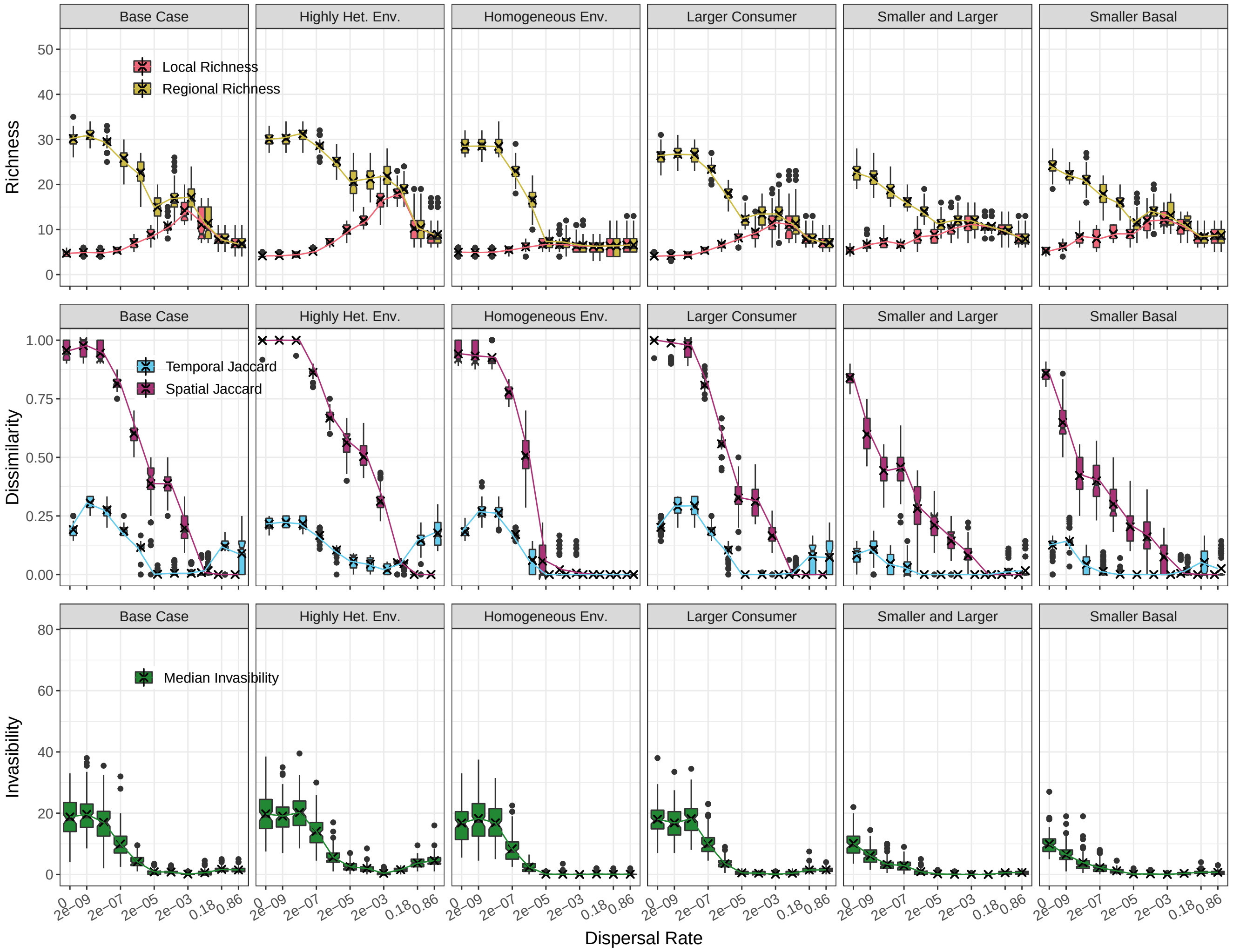


**Fig. S12.** As in Figure S11, but instead, columns represent the variations in the Pool & Noise structure. From left to right, the columns refer to the base case, to the highly heterogeneous environments case (i.e., doubled coefficient of variation of the interaction matrices), to the homogeneous environments case (i.e., no noise in the interaction matrix), to larger consumer species (i.e., an increase of 1 to the maximum logarithmic body size), to larger consumer species and smaller basal species (i.e., a decrease of 1 to the minimum logarithmic body size), and to smaller basal species. As discussed in Section 4, the largest differences in the patterns are observed with the homogeneous patch environments case, in which communities homogenise for lower dispersal values than otherwise observed. Axes are chosen to match Figure S11.

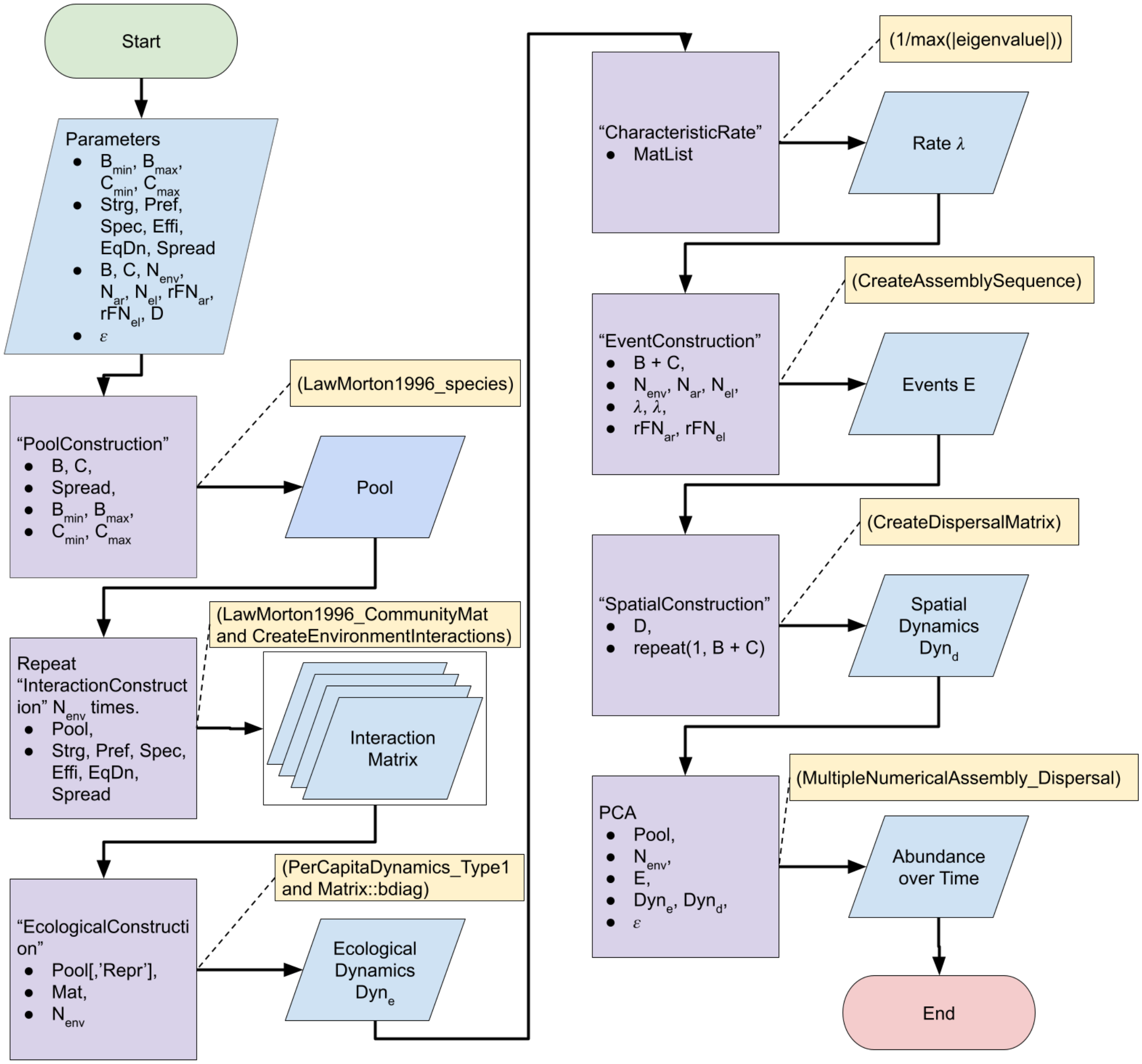


**Fig. S13.** Diagram of an Example Simulation Run. Arrows represent flow through the example run, while dashed lines to rectangles are used to notate the actual functions used in the R code. Parallelograms represent inputs and outputs to the process while squares represent algorithm steps. Algorithm steps are described in more detail in Algorithms 1 - 7.
